# scCody distinguishes aging-aligned from divergent disease-associated cellular remodeling

**DOI:** 10.64898/2026.11.10.765622

**Authors:** Jiramet Kinchagawat, Jinmiao Chen, Sai Zhang, Xiaotao Shen

## Abstract

Biological aging clocks summarize molecular changes as a single age estimate, but cannot distinguish disease-associated changes that follow healthy aging from those that deviate from it. Here we introduce scCody, a generative framework that learns age-dependent cellular population dynamics from cross-sectional single-cell cohorts using population-level optimal transport and endpoint-consistency regularization. scCody accurately reconstructed cellular populations at unseen ages, recovered age-associated transcriptional changes more effectively than existing generative dynamics models, and enabled donor-age inference without direct age-regression training. Using the learned healthy aging reference, scCody decomposed disease-associated cellular displacement into aging-aligned and orthogonal components. In Alzheimer’s disease, selected neuronal populations showed substantial aging-aligned remodeling alongside shared orthogonal changes associated with established pathology signatures. In contrast, COVID-19 exhibited weaker aging alignment and more pronounced orthogonal immune remodeling. Further decomposition revealed patient-level interferon variation associated with symptom duration and increased transcriptional heterogeneity in CD16 monocytes. This framework provides an extension of biological age assessment beyond a single age estimate, distinguishing cellular changes that resemble healthy aging from those that depart from it.

## Introduction

Population-scale single-cell RNA sequencing (scRNA-seq) cohorts now profile hundreds to thousands of individuals across the adult lifespan, providing an opportunity to characterize healthy human aging at cellular resolution^1–3^. These datasets can reveal how cellular states and their relative abundances change with age, how aging trajectories differ across cell types^4^, and how disease-associated cellular states relate to those observed during healthy aging. Establishing such a reference is particularly important given the heterogeneity of aging across individuals, tissues, and cellular populations.

Biological aging is commonly assessed using supervised aging clocks, which predict chronological age from molecular profiles, with deviations between predicted and chronological age often interpreted as indicators of biological aging^5–8^. Recent single-cell aging clocks have extended this approach to individual cellular lineages, revealing disease-associated age shifts and heterogeneity in cellular aging^9,10^. However, supervised aging clocks have an important limitation when applied to disease. These models are trained to predict chronological age rather than explicitly model healthy aging trajectories. Consequently, disease-associated transcriptional changes that deviate from healthy aging may nevertheless influence predicted age. A positive age gap therefore does not necessarily indicate accelerated healthy aging, while a near-zero age gap does not exclude substantial disease-associated dysregulation. With another study using clinical records to disentangle healthy aging from chronic disease risk^11^, the same principle should also be applied to molecular clocks. Given that molecular aging involves both coordinated and stochastic changes^12,13^, a single age estimate cannot distinguish disease-associated changes that follow healthy aging from those that deviate from it, highlighting the need for an explicit reference of healthy cellular aging.

A complementary strategy is to model age-dependent distributions of cellular states using generative approaches, which can capture changes in gene expression, cellular heterogeneity, and cell-state abundance^14,15^. Optimal transport and continuous-time models have enabled the modeling of cellular dynamics without known cell-to-cell correspondence, particularly in developmental and perturbation settings^16–21^. However, human aging cohorts present a distinct challenge: cellular populations at different ages are sampled from different individuals, and observed differences reflect not only aging but also donor-specific biological variation. Learning healthy aging dynamics from such cohorts therefore requires a framework that captures coherent age-dependent population changes while minimizing dependence on the particular donor populations used as starting points.

Here we present scCody (single-cell Cohort Dynamics), a generative framework for learning age-conditioned cellular population dynamics from cross-sectional single-cell cohorts. Building on FlowMap^22^, scCody learns cellular transport between ages through population-level distribution matching without requiring cell-to-cell correspondence. We introduce an endpoint-consistency regularizer designed to encourage populations transported from different source ages to converge toward consistent target-age distributions. The resulting generative reference enables reconstruction of cellular populations at unseen ages and inference of donor age through trajectory projection without direct age-regression training. Unlike a supervised clock, the resulting trajectory-derived age is determined by the donor’s position relative to a learned healthy aging trajectory, providing an estimate of where its cellular population lies along the patterns observed during healthy aging.

Disease-associated cellular changes can occur at multiple levels, reflecting both shared disease responses and heterogeneity between and within individuals. Previous hierarchical models have distinguished between-individual and within-individual variation in multi-subject scRNA-seq data^23,24^, while more recent approaches have shown that these effects can differ across cell types^25^ or cellular states^15^. Building on these concepts, scCody characterizes disease-associated displacement relative to the learned healthy aging reference. It first separates displacement into an aging-aligned component, representing changes along or against the healthy aging direction, and an orthogonal component, representing changes that deviate from this direction. The orthogonal component is further decomposed into a shared shift across patients, between-patient variation, and within-patient variation. Together, these decompositions characterize both how disease-associated changes relate to healthy aging and how they are organized across patients and individual cells.

We evaluated scCody through population reconstruction, age inference, and transcriptional aging-program analyses, and applied it to Alzheimer’s disease (AD) and COVID-19. We found substantial aging-aligned remodeling in selected brain cell populations in AD, whereas COVID-19-associated immune changes were characterized by pronounced orthogonal displacement. Further decomposition revealed shared pathology-associated neuronal remodeling in AD and distinct interferon-associated and monocyte responses in COVID-19. This framework extends biological aging analysis beyond a single age estimate.

## Results

### Overview of scCody

scCody is a generative framework designed to learn age-dependent cellular population dynamics from cross-sectional single-cell cohorts. Unlike longitudinal studies, cross-sectional cohorts sample different individuals at different ages, making it chalzzlenging to distinguish shared aging-related changes from donor-specific variation (Fig. 1a). Rather than directly predicting chronological age, scCody learns how cellular population distributions change across the lifespan, establishing a generative reference of healthy aging within the broader cellular state space (Fig. 1b). We used the Asian Immune Diversity Atlas (AIDA)^2^ as the reference cohort and selected 2,000 highly variable genes (HVGs) for model training (Fig. 1c, Supplementary Fig. 1, Supplementary table 1 and Methods).

**Figure 1.**
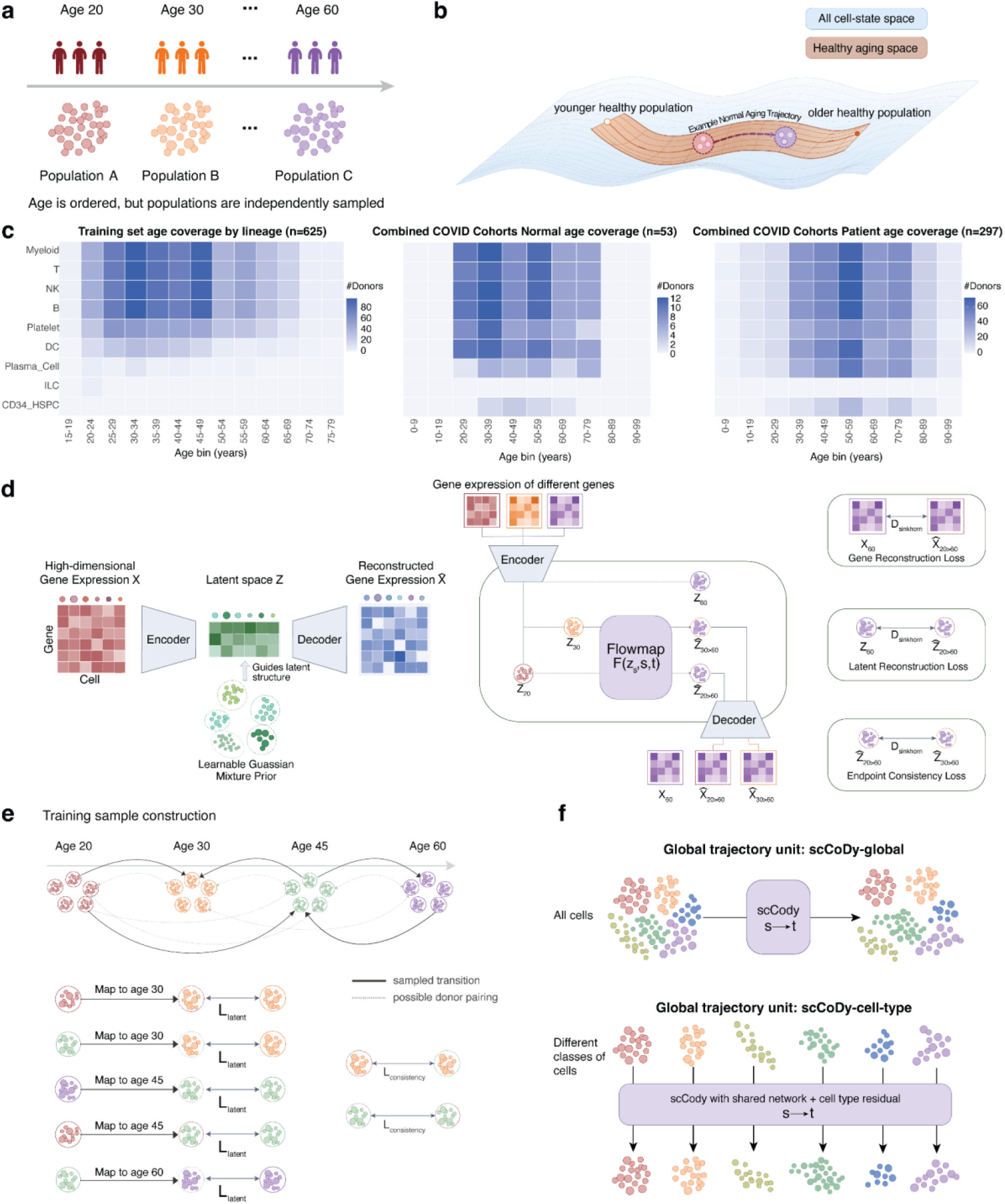
Overview of scCody for modeling age-dependent cellular population dynamics from cross-sectional single-cell cohorts. (a) Cross-sectional aging cohorts comprise independently sampled donors and cellular populations ordered by chronological age, without longitudinal correspondence between individuals or cells. (b) Conceptual representation of healthy aging within the broader cellular state space. Healthy populations occupy heterogeneous, age-dependent distributions that define a reference for modeling age-associated changes and assessing disease-associated deviations. (c) Donor coverage across age bins and major cellular lineages in the healthy training cohort (left) and the healthy control (middle) and patient (right) groups of the combined COVID-19 cohorts. (d) scCody architecture and training objectives. An encoder-decoder maps gene-expression profiles into a latent space structured by a learnable Gaussian-mixture prior. An age-conditioned flow map transports latent cellular states from source age to target age, and a decoder generates target-age expression profiles. Training incorporates gene-expression reconstruction, latent-space population matching, and endpoint consistency. (e) Construction of population-level training examples. Source and target donor populations are sampled across ages for younger-to-older and older-to-younger transport. Transported populations are compared with observed target-age populations using Sinkhorn-based optimal transport. Endpoint consistency additionally encourages agreement between populations transported from different source ages to a common target age. (f) Complementary modeling formulations. scCody-G transports pooled populations across cell types. scCody-CT combines a shared network with a cell-type-specific residual component to model transcriptional changes within cellular lineages.

scCody combines a structured latent representation with age-conditioned cellular transport (Fig. 1d and Methods). An encoder maps single-cell gene-expression profiles into a lower-dimensional latent space that accommodates heterogeneous cellular states. An age-conditioned flow map then transports these latent states from a source age to a specified target age, and a decoder reconstructs the corresponding gene-expression profiles. Because cells observed at different ages lack known cell-to-cell correspondence, scCody learns these transformations by matching generated and observed target-age populations (Methods).

However, population-level matching alone may still produce source-dependent predictions, as different starting populations can yield distinct endpoints, particularly in the presence of donor heterogeneity. To address this challenge, scCody introduces an endpoint-consistency regularizer tailored to cross-sectional cohort data (Fig. 1e and Methods). During training, populations sampled from different source ages are independently transported toward a common target age. In addition to matching each generated population to the observed target distribution, the regularizer penalizes disagreement among the independently generated endpoints. This soft population-level constraint reduces source dependence while preserving cellular heterogeneity.

We implemented two complementary formulations to characterize aging at different cellular resolutions (Fig. 1f). scCody-G (global) models pooled cellular populations, capturing age-dependent changes in both cell-type composition and transcriptional states. scCody-CT (cell type) extends the shared transport network with cell-type-specific residual components, allowing distinct age-dependent transcriptional changes across cellular lineages.

### scCody reconstructs cellular populations at unseen ages

To determine whether scCody captures age-dependent cellular dynamics beyond the populations observed during training, we evaluated its ability to reconstruct cellular populations at unseen ages. We established a benchmark using a healthy peripheral blood mononuclear cell (PBMC) single-cell RNA-sequencing dataset comprising 625 donors and approximately 1.27 million cells. All cells from donors aged 21, 31, 41, 51, 61, and 68 years were excluded from model training, leaving 554 donors to establish the healthy aging reference (Fig. 2a and Methods). We then used the learned dynamics to generate populations at these six held-out ages and compared the performance of scCody-G and scCody-CT with three continuous-time generative models: scNODE^18^, scDiffEq^19^, and ARTEMIS^20^ (Methods and Supplementary Note 1).

**Figure 2.**
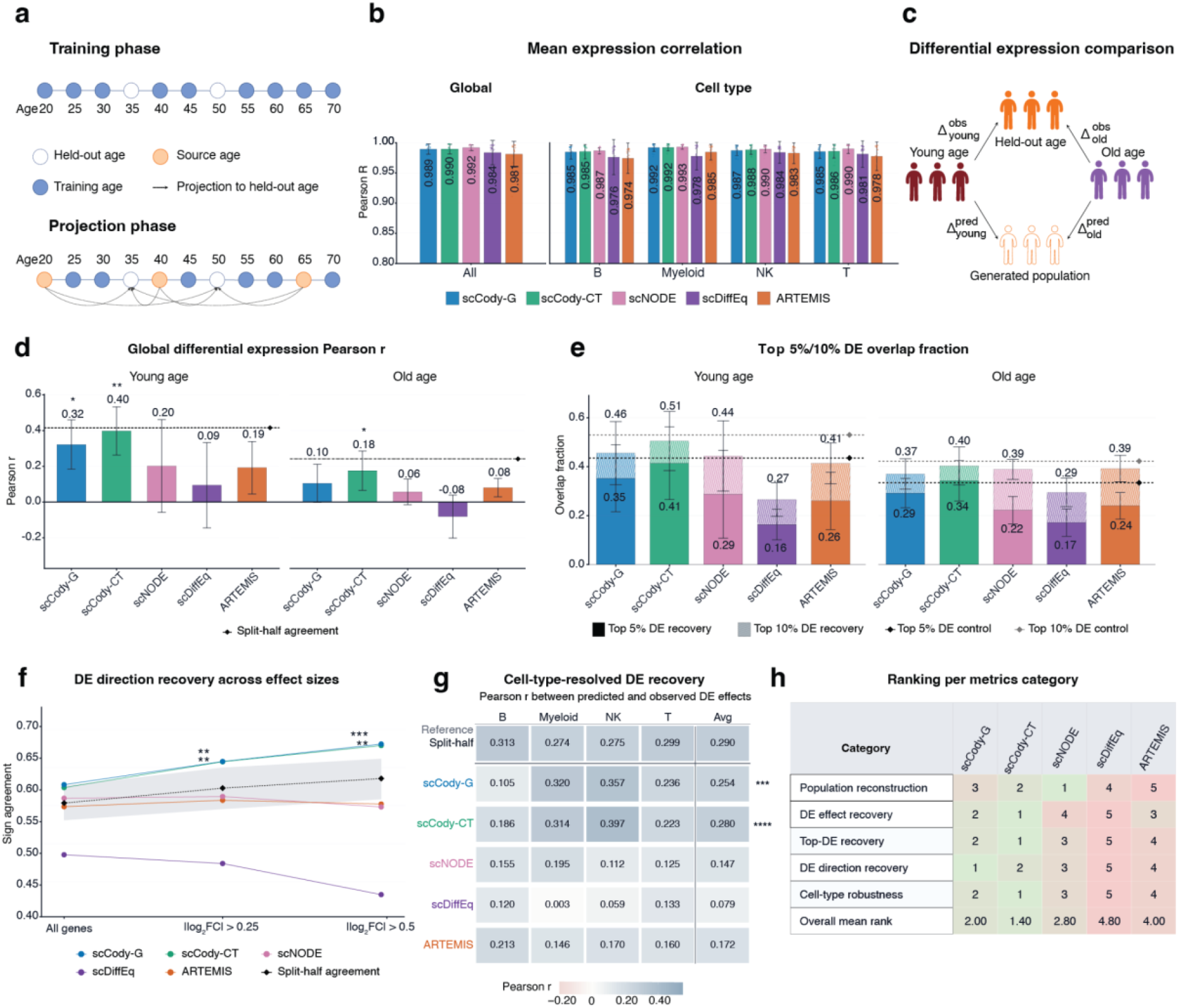
Benchmarking reconstruction of cellular populations and age-associated transcriptional changes at held-out ages. (a) Held-out-age evaluation using a healthy PBMC single-cell RNA-sequencing dataset comprising 625 donors and approximately 1.27 million cells. All cells from donors within held-out ages were excluded from training (Ages shown are illustrative; see Methods for held-out ages). Models trained on the remaining ages generated populations at the held-out ages from observed source populations. (b) Pearson correlations across genes between generated and observed mean-expression profiles, evaluated across all cells and within B, myeloid, natural killer (NK), and T cells. (c) Differential-expression (DE) evaluation scheme. Generated and observed populations at each held-out age were each compared with younger and older reference populations, yielding predicted and observed DE effects for the two reference comparisons. (d) Pearson correlations between predicted and observed gene-wise log_2_ fold changes across the pooled population for younger-reference (left) and older-reference (right) comparisons. (e) Overlap fractions between predicted and observed sets of the top 5% and top 10% of DEs for younger-reference (left) and older-reference (right) comparisons. (f) Concordance between predicted and observed directions of expression change across genes with nonzero effects and at absolute log2 fold-change thresholds of 0.25 and 0.5. (g) Cell-type-resolved DE recovery, measured by Pearson correlation between predicted and observed gene-wise log2 fold changes. (h) Method rankings across six evaluation categories and the overall mean rank.

All models achieved high correlations between generated and observed mean gene expression, with global Pearson correlations exceeding 0.97 and similarly high agreement within B, myeloid, natural killer (NK), and T cells (Fig. 2b and Supplementary Fig. 2a). scNODE achieved the highest global correlation of 0.992, followed by scCody-CT and scCody-G at 0.990 and 0.989, respectively. However, the consistently high correlations across models indicated that mean-expression agreement alone was insufficient to distinguish their ability to recover age-associated transcriptional changes.

We therefore examined whether the generated populations reproduced quantitative differential-expression (DE) effects between ages. For each held-out age, the generated and observed populations were separately compared with the same younger and older reference populations, yielding predicted and observed gene-wise log_2_ fold changes for younger-reference (DE_young_) and older-reference (DE_old_) comparisons (Fig. 2c and Methods). To contextualize model performance, we also split the observed cells into two halves and independently estimated the corresponding DE effects, providing an empirical reference for reproducibility (Methods). scCody-CT achieved correlations of 0.40 for DE_young_ and 0.18 for DE_old_, while scCody-G got 0.32 and 0.10, compared with 0.20 and 0.06 for scNODE, 0.09 and - 0.08 for scDiffEq, and 0.19 and 0.08 for ARTEMIS, respectively (Fig. 2d and Supplementary Data 1). The correlations achieved by scCody-CT approached the empirical split-half agreement for both DE_young_ and DE_old_ (Supplementary Fig. 2b). These results indicate that scCody more accurately recovers the magnitude and relative patterns of age-associated transcriptional changes than the other evaluated generative models.

scCody also recovered a larger fraction of the top 10% of age-associated DE genes than the comparison models, although its advantage over scNODE was smaller among the top 5% (Fig. 2e). Gene recovery was evaluated by ranking predicted and observed DE effects by magnitude and calculating the overlap between their top-ranked gene sets (Methods). Directional agreement generally increased with the absolute log_2_ fold-change threshold, with scCody showing higher DE sign concordance than the other generative models across genes with nonzero effects and at the evaluated effect-size thresholds (Fig. 2f).

scCody-CT recovered the largest fraction of top-ranked age-associated DE genes, with overlaps of 0.41 and 0.51 for the top 5% and 10% of genes in younger-reference comparisons, although its advantage over scNODE and ARTEMIS was modest (Fig. 2e). Gene recovery was evaluated by ranking predicted and observed DE effects by magnitude and calculating the overlap between their top-ranked gene sets (Methods). Directional agreement generally increased with the absolute log_2_ fold-change threshold. Across all genes with nonzero effects, directional agreement was comparable between scCody and scNODE (57.6-60.8% and 57.7-58.7%, respectively). However, agreement for scCody increased with the absolute log_2_ fold-change threshold, reaching 67.2% and 68.9% among genes with |log_2_FC| > 0.5 in younger- and older-reference comparisons, whereas scNODE and ARTEMIS remained between 56.5% and 59.5%, and scDiffEq declined to 43.4% and 50.5% (Fig. 2f). These results indicate that scCody most consistently recovers the direction of larger age-associated transcriptional changes.

We then examined whether these improvements extended to individual immune lineages ( Methods). Across B, myeloid, NK, and T cells, scCody-G and scCody-CT achieved mean Pearson correlations of 0.254 and 0.280, respectively, compared with 0.147 for scNODE, 0.079 for scDiffEq, and 0.172 for ARTEMIS (Fig. 2g). Both scCody implementations outperformed the comparison models in myeloid, NK and T cells and on average, with scCody-CT approaching split-half agreement (0.280 vs 0.290). These findings indicate that scCody captures age-associated transcriptional changes within individual immune lineages rather than relying solely on differences in cell-type composition.

Finally, we summarized model performance across six evaluation categories (Fig. 2h). scCody-CT ranked first overall across all six categories, while scCody-G ranked second overall, with mean ranks of 1.40 and 2.00, respectively. Although these rankings do not account for differences in effect magnitude across metrics, they were consistent with the individual benchmark results.

### Learned aging trajectories recover donor age and age-related immune changes

Given that scCody can reconstruct cellular populations at unseen ages, we next asked whether the learned dynamics could be used to infer the age of an unseen donor. scCody generates healthy reference populations across candidate ages and identifies the age at which the generated population most closely matches the observed donor population. Specifically, for each held-out donor, we transported healthy reference populations across candidate ages and calculated the Sinkhorn distance^26^ between each generated population and the observed donor population (Fig. 3a and Methods). The candidate age minimizing this distance was taken as the trajectory-derived age estimate. To reduce dependence on the starting population, we repeated this procedure using multiple healthy reference populations and aggregated their estimates. Representative donors aged 41 and 61 years were assigned ages of 39.2 and 61.0 years, respectively (Fig. 3b).

**Figure 3.**
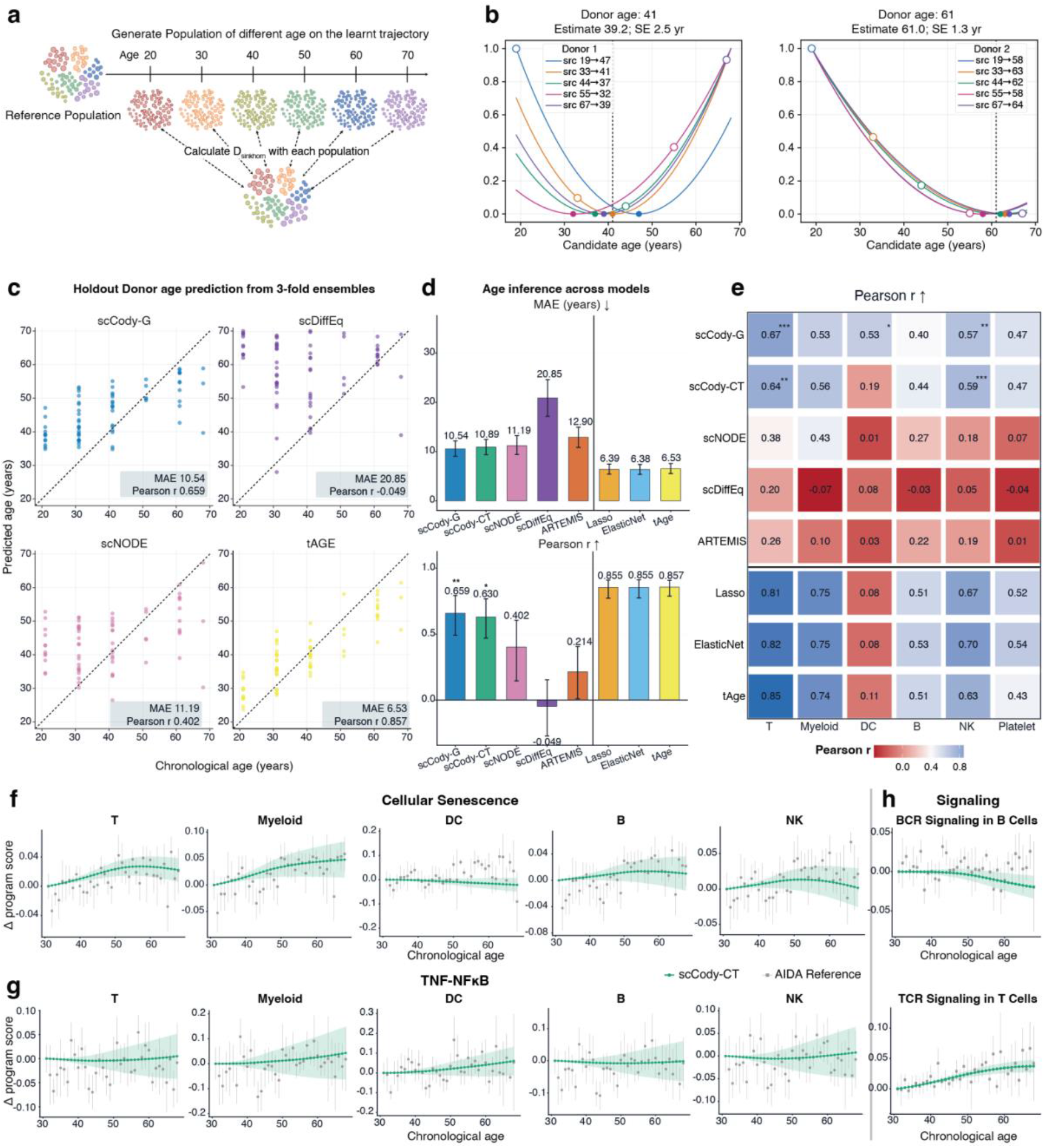
Generative aging trajectories enable donor-age inference and recover age-associated transcriptional programs. (a) Overview of generative trajectory projection. A healthy reference population is transported across candidate ages using the learned scCody dynamics. Generated populations are compared with an unseen query population using Sinkhorn distance, and the candidate age minimizing this distance provides a source-specific age estimate. Estimates obtained from multiple reference populations are aggregated to infer donor age. (b) Source-specific distance curves for two example donors aged 41 years (left) and 61 years (right). Colors indicate different reference populations; open circles mark source ages, and filled circles mark distance minima. Legend entries indicate source age followed by the inferred age. Vertical dotted lines indicate chronological age. Mean age estimates and their standard errors (SEs) across source trajectories are shown above each plot. (c) Predicted versus chronological age in held-out healthy donors using three-fold model ensembles, shown for scCody-CT, scDiffEq, ARTEMIS and tAge. Each point represents a donor, and dotted lines indicate identity. (d) Comparison of donor-age inference across scCody-G, scCody-CT, scNODE, scDiffEq, ARTEMIS and supervised age-prediction models (Lasso, ElasticNet and tAge), evaluated using MAE (top; lower is better) and Pearson correlation (bottom; higher is better). The vertical line separates dynamic generative models from supervised prediction models. (e) Cell-type-resolved age inference. Heatmap entries show Pearson correlations between predicted and chronological age within B, myeloid, natural killer (NK), platelet, and T-cell populations. Colors indicate correlation magnitude and direction. (f-h) Age-related pathway changes along scCody-CT trajectories. Starting from the held-out age-31 population, populations were generated at yearly intervals and scored for senescence-(f) and TNF–NFκB-associated (g) expression across immune cell types, B-cell receptor (BCR)-associated expression in B cells, and T-cell receptor (TCR)-associated expression in T cells (h). Green trajectories show scCody-CT-generated populations with 95% bootstrap confidence intervals, and gray values show the corresponding scores in observed AIDA populations across age.

We evaluated trajectory-derived age estimates in held-out healthy donors using three-fold model ensembles (Fig. 3c,d, Supplementary Fig. 3, Supplementary Data 2 and Methods). Among the evaluated generative dynamics models, scCody-G achieved the lowest mean absolute error (MAE) of 10.54 years and a Pearson correlation of 0.659. scCody-CT achieved an MAE of 10.89 years and a Pearson correlation of 0.63, whereas scNODE, scDiffEq, and ARTEMIS yielded MAEs ranging from 11.19 to 20.85 years (Fig. 3d). Thus, scCody recovered chronological age more accurately than the other generative dynamics models despite not being trained with an explicit age-regression objective. Supervised age-prediction methods, including Lasso and ElasticNet, which are popular choices across several published literature^9,27^, and tAge^28^ achieved lower MAEs and higher Pearson correlations (Fig. 3d and Methods), confirming their advantage in direct chronological age prediction. However, scCody also provides a generative reference for examining how cellular populations change across age.

Similar to the cell-type-resolved DE analysis, scCody also recovered age-associated information within individual immune lineages (Fig. 3e), although performance varied considerably more across cell types. For scCody-CT, correlations between predicted and chronological age were highest in T cells (r = 0.64), followed by NK cells (r = 0.59), myeloid cells (r = 0.56), and platelets (r = 0.47). Correlations were weaker in B cells (r = 0.44) and dendritic cells (r = 0.19). scCody-G showed a similar pattern, with correlations ranging from 0.40 to 0.67 across the four major immune lineages, but a stronger association in dendritic cells (r = 0.53). The other generative dynamics models generally showed limited within-lineage age recovery, particularly scDiffEq, which models dynamics in a principal-component space, possibly because age-associated variation is weakly represented among the leading components. Supervised models performed better in several cell types, particularly T, myeloid, and NK cells. Across scCody-CT and supervised models, the per-cell-type pattern was broadly similar, with stronger age prediction in T cells and weaker performance in dendritic cells. This suggests that some immune lineages may contain less age-related information in their transcription or be more inherently difficult to model.

We assessed the contributions of the Gaussian-mixture prior, endpoint consistency, and latent optimal-transport regularization through ablation analyses (Supplementary Table 2). Replacing the Gaussian-mixture prior with a standard VAE prior produced the largest increase in held-out Sinkhorn divergence, and removing either regularizer also increased it in both formulations. In the three-fold validation setting, removing endpoint consistency additionally reduced the correlation between inferred and chronological age in both formulations (r = 0.716 vs 0.591 for scCody-G and 0.682 vs 0.599 for scCody-CT), although this effect was not observed on the external held-out ages.

### scCody recapitulates cell-type-specific transcriptional programs of aging

We next examined whether scCody reproduced established transcriptional programs of immune aging. Starting from held-out donors aged 31 years, we generated populations at yearly intervals and compared their program scores with those observed across ages in AIDA (Fig. 3f-h, Supplementary Data 3 and Methods). Generated trajectories followed the observed increase in senescence-associated expression in T and myeloid cells, with the largest increase in myeloid cells (Fig. 3f), and TCR-associated expression increased with age in both generated and observed T cells (Fig. 3h). The T-cell increase is consistent with the age-related accumulation of p16INK4a transcripts in human peripheral blood T cells^29^.

Inflammatory programs showed pronounced cell-type specificity. Across the young-older window, TNF–NFκB-associated expression increased in dendritic and myeloid cells, whereas T, B and NK cells remained comparatively stable (Fig. 3g). This pattern agrees with single-cell comparisons of young and older donors reporting the accumulation of inflammatory monocytes and age-associated dendritic cells^30^, and with transcriptional evidence of a semi-activated state with elevated basal NF-κB activity in circulating dendritic cells from older individuals^31,32^.

Several trajectories were non-monotonic at older ages. Immune-aging programs have largely been defined by contrasting younger and older donors, such as adults aged 25-35 and 55-65 years^33^, and across this window, senescence-associated expression increased in T, myeloid, B, and NK cells, consistent with the reported direction of change. The generated T-cell trajectory also resembled the nonlinear pattern reported in age-susceptible T-cell subsets of the follow-up cohort of Gong *et al*., which increased with age before plateauing, although this plateau was not observed in their two other cohorts^1,34^. This disagreement may partly reflect the sensitivity of trajectories to each cohort’s age distribution^35^ and the sparse support for older ages in our training set. Generated B cells further showed a decline in BCR-associated expression, compatible with reduced BCR responsiveness in aging B cells^36^. Together, scCody reproduced the direction and cell-type specificity of established immune-aging programs across adulthood.

### scCody distinguishes aging-aligned from orthogonal disease-associated changes

Disease-associated transcriptional changes may follow the direction of healthy aging, deviate from it, or involve both processes. We hypothesized that these contributions differ across disease contexts and examined whether generative and supervised aging models produce distinct age-shift estimates in two contrasting diseases: AD, a chronic neurodegenerative disorder strongly associated with aging, and COVID-19, an acute infection characterized by immune system perturbation.

We analyzed single-nucleus RNA-seq data from 212 donors with AD in the PsychAD cohort in MSSM subset^37^ and peripheral-blood single-cell RNA-seq data from 230 patients with COVID-19 across three independent cohorts^38–40^. For AD, we trained a separate scCody model on healthy donors and evaluated it in held-out healthy donors (Supplementary Fig. 4). Disease-associated age shifts were estimated relative to sex- and age-matched healthy donors from the corresponding cohorts (Methods). In AD, scCody estimated older cellular ages in most cell types, with the largest shifts in excitatory neurons and astrocytes (Fig. 4a). ElasticNet, a supervised aging clock trained on the same healthy reference, produced broadly similar shifts in several populations, including astrocytes, immune cells, and oligodendrocyte progenitor cells, but showed little acceleration in excitatory neurons. In contrast, the two approaches diverged substantially in COVID-19. scCody estimated relatively small age shifts across the three cohorts, whereas ElasticNet produced larger and more variable shifts, reaching approximately 10 years in myeloid cells and platelets (Fig. 4a and Supplementary Fig. 5a and Supplementary Data 4). Donor-level concordance between the two approaches was moderate in AD (Lin’s concordance correlation coefficient, CCC = 0.52, mean cell-type CCC = 0.50) but lower in COVID-19 (CCC = 0.29, mean cell-type CCC = 0.36) (Supplementary Fig. 5b,c).

**Figure 4.**
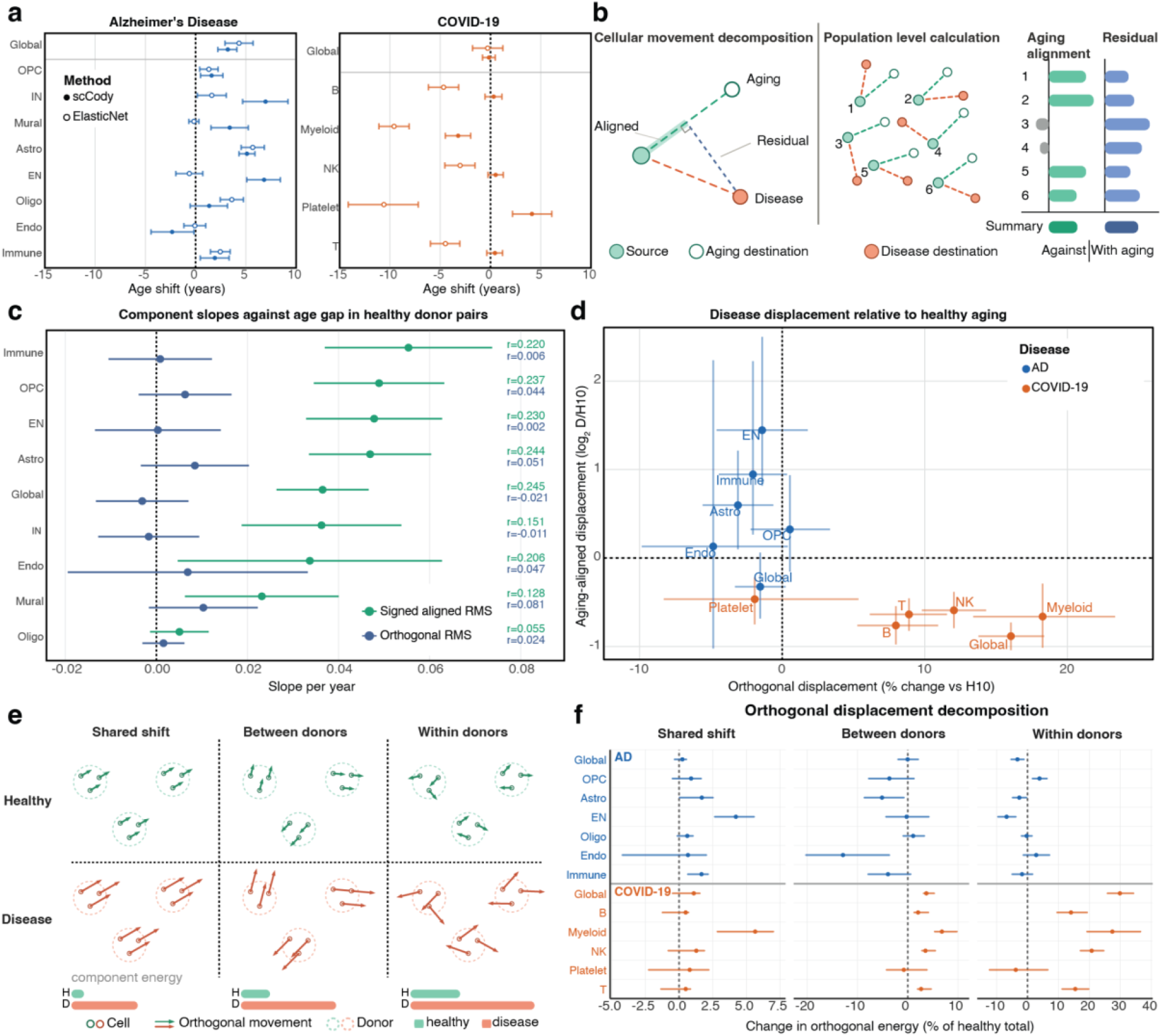
Decomposition of disease-associated cellular displacement relative to healthy aging. (a) Age shift in AD (left) and COVID-19 (right), estimated by scCody (filled circles) and ElasticNet (open circles). Age shift is the inferred minus chronological age gap difference between disease donors and age-matched healthy donors within a 5-year window, using donors not used for training. COVID-19 estimates are pooled across three cohorts (Ren, COMBAT, and Stephenson). ElasticNet was trained on the same healthy reference as scCody. Points show estimates and bars show 95% bootstrap confidence intervals. (b) Schematic of the displacement decomposition. For each source cell, the aging displacement is given by the learned scCody dynamics and the disease displacement by optimal-transport matching to disease cells. Disease displacement is decomposed into a component aligned with the aging direction and an orthogonal component. Each is summarized across cells as a signed aligned root mean square (RMS) and an orthogonal RMS. The same quantities are computed for healthy donor pairs separated by 10 years of age, which serve as the reference. (c) Validation of the aging direction in healthy donor pairs. Slope per year of age gap for signed aligned RMS (green) and orthogonal RMS (blue), with Pearson correlations with age gap shown on the right. Bars show 95% donor-level bootstrap confidence intervals. The aligned component increases with age gap in most cell types, while the orthogonal displacement shows substantially weaker age dependence. (d) Disease displacement relative to 10 years of healthy aging. The y-axis shows the log_2_ ratio of signed aligned RMS in disease relative to the healthy reference, and the x-axis the percentage change in orthogonal RMS relative to the same reference. Each point is a cell type in AD (blue) or pooled COVID-19 (orange). Bars show 95% bootstrap confidence intervals, and arrows mark intervals extending beyond the plotted range. (e) Schematic of the orthogonal decomposition (illustrative, not data). Orthogonal displacements are partitioned into a component shared across donors, variation between donor means, and variation within donors. The partition is computed in the same way for healthy donor pairs separated by 10 years (green) and for disease pairs (red). Bars below each panel indicate the energy of each component. (f) Change in each orthogonal component in disease relative to the healthy reference, expressed as a percentage of the healthy total orthogonal energy, so that the three components sum to the change in total orthogonal energy. Points show estimates and bars show 95% donor-level bootstrap confidence intervals. Dashed lines mark no change.

To directly characterize these changes, we used the age-conditioned dynamics learned by scCody to decompose disease-associated cellular displacement (Fig. 4b and Methods). For each cell, scCody defined a healthy aging displacement, while optimal-transport matching between healthy and disease populations estimated the corresponding disease-associated displacement. The latter was separated into an aging-aligned component, representing changes along or against the healthy aging direction, and an orthogonal component, representing changes perpendicular to that direction. These components were quantified using signed aligned root mean square (signed aligned RMS) and orthogonal root mean square (orthogonal RMS), respectively.

We first validated this decomposition using healthy donors from the PsychAD cohort, for whom exact chronological ages were available. Cellular displacements between healthy donor pairs spanning different age differences were decomposed using the learned aging direction (Methods). Aging-aligned displacement increased with chronological age difference across most brain cell populations, whereas orthogonal displacement showed substantially weaker age dependence (Fig. 4c). This pattern was evident globally and within several cell types, although the aging-aligned association was weaker in oligodendrocytes. We therefore used the displacement between healthy donors separated by ten years as an empirical reference for evaluating disease-associated remodeling.

Relative to this reference, AD and COVID-19 exhibited markedly different displacement patterns (Fig. 4d and Supplementary Data 5). In AD, aging-aligned displacement exceeded the ten-year healthy-aging reference in excitatory neurons, immune cells, and astrocytes, reaching 2.7-fold the reference in excitatory neurons. In contrast, COVID-19 showed weaker aging-aligned displacement across all examined cell types, as well as globally. Meanwhile, orthogonal displacement increased across all contrasts with the exception of platelets, whose orthogonal displacement decreased. These findings help explain the age-shift differences observed in Fig. 4a. In several AD populations, broadly similar age-shift estimates from scCody and ElasticNet coincided with substantial aging-aligned displacement. In COVID-19, the models yielded divergent age-shift estimates, while the displacement analysis revealed weaker aging alignment and pronounced orthogonal remodeling (Fig. 4a,d). Age-shift estimates alone therefore do not establish the extent to which disease-associated cellular changes follow healthy aging.

### scCody resolves patient- and cell-level heterogeneity in orthogonal disease-associated remodeling

The cell-level displacements derived from optimal-transport coupling allowed us to further examine how orthogonal disease-associated changes were distributed across patients and cells. We decomposed orthogonal displacement into three components: a shared shift across patients, variation between patient means, and variation among cells within individual patients (Fig. 4e). Each component was compared with its corresponding value in healthy donor pairs separated by ten years, providing an empirical reference for assessing disease-associated changes (Fig. 4f and Methods).

In AD, shared orthogonal displacement increased across several brain cell populations, most prominently in excitatory neurons, indicating a common disease-associated transcriptional shift beyond the healthy aging direction (Fig. 4f). In contrast, between-patient variation decreased in endothelial cells and astrocytes, while within-patient variation decreased in excitatory neurons. The latter finding is consistent with the reported selective loss of vulnerable neuronal subpopulations in AD^41^, although changes in neuronal population structure may also contribute. Thus, AD-associated orthogonal remodeling was characterized by a prominent shared neuronal shift alongside selective reductions in patient- and cell-level heterogeneity.

COVID-19 showed a different pattern, with shared displacement increased mainly in myeloid cells, while between-patient and within-patient variation increased across most cell types, with within-patient variation increasing most, particularly in myeloid cells (Fig. 4g). These findings indicate that COVID-19-associated orthogonal remodeling was dominated by increased cellular heterogeneity within individual patients, contrasting with the prominent shared neuronal displacement observed in AD.

### Disease-associated displacement reveals neuronal remodeling in AD and heterogeneous myeloid responses in COVID-19

The orthogonal decomposition highlighted a major shared displacement in AD excitatory neurons and a marked increase in within-patient residual energy in COVID-19 myeloid cells (Fig. 4f). To identify the transcriptional programs associated with these changes, we examined the decoded displacement components in these populations (Methods).

In AD excitatory neurons, the decoded displacements were concordant with pathology-associated transcriptional signatures identified in an independent cohort^42,43^. Genes previously reported to increase with AD pathology showed positive enrichment relative to ten years of healthy aging in both the aging-aligned and shared components (z = 8.2 and 6.1, respectively; Fig. 5a, Supplementary Data 6 and Methods). Conversely, pathology-associated downregulated genes showed negative enrichment in both components (z = -6.5 and -6.6). Synaptic-signaling genes exhibited a similar distribution across the aging-aligned and shared components (z = 4.3 and 2.7). The decomposition thus attributed these AD pathology-associated transcriptional changes to both aging-aligned and shared orthogonal displacement.

**Figure 5.**
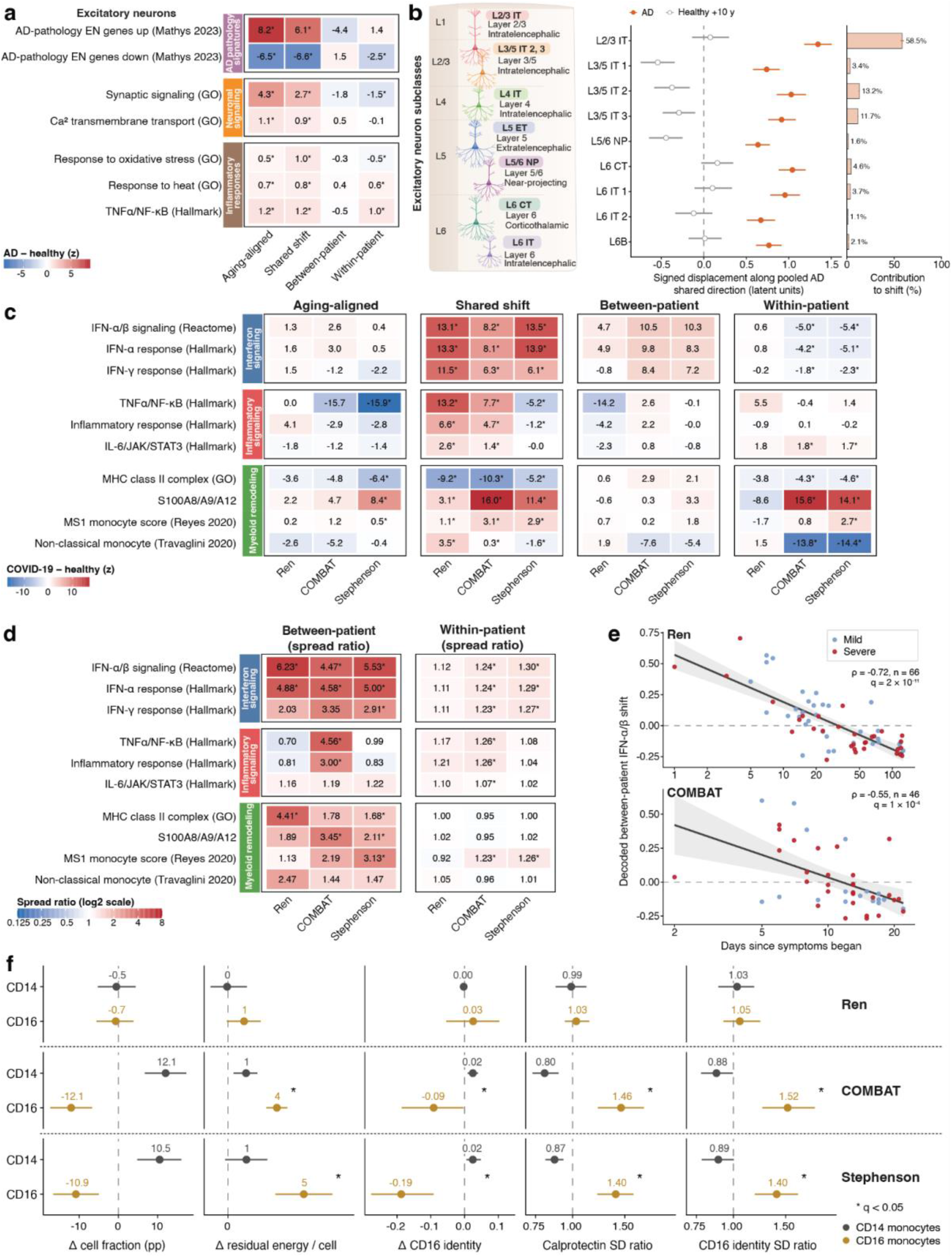
Molecular and cellular characterization of disease-associated displacement in AD and COVID-**19**. (a) Enrichment of decoded gene-program changes across aging-aligned, shared, between-patient and within-patient components in AD excitatory neurons. Values are z-scores of AD-associated changes relative to a ten-year healthy-aging reference, standardized against 2,000 size-matched random gene sets. (b) Shared displacement across AD excitatory-neuron subclasses. Left, the illustration of excitatory-neuron subclasses. Middle, signed projections onto the pooled shared direction in AD (filled circles) and healthy controls (open circles). Right, subclass contributions to the population-level shared shift, weighted by cell fraction. Error bars indicate 95% donor-bootstrap confidence intervals. (c) Gene-program enrichment across the same four displacement components in COVID-19 myeloid cells from Ren, COMBAT, and Stephenson, calculated as in a. Colored bars indicate biological program groups. (d) Between-patient and within-patient dispersion of decoded myeloid programs, expressed as COVID-19/healthy standard-deviation ratios. Between-patient dispersion is calculated across donor means; within-patient dispersion is the mean within-donor standard deviation. (e) Decoded between-patient IFN-α/β response versus days since symptom onset in Ren (n = 66) and COMBAT (n = 46). Points represent patients colored by severity; lines show least-squares fits against log-transformed days with 95% confidence bands. Spearman correlations are indicated. (f) CD14/CD16 monocyte comparison across COVID-19 cohorts. From left to right, changes in cell fraction, within-patient residual energy per cell, CD16-identity program score, and within-patient dispersion of calprotectin and CD16-identity programs. Points show estimates with 95% donor-bootstrap confidence intervals; within-patient analyses use same-subtype matches. The healthy reference consists of control populations separated by approximately ten years, evaluated using the same decomposition. Asterisks indicate Benjamini-Hochberg-adjusted q < 0.05 (a,c, program changes; d, increased dispersion; f, direct CD14-CD16 contrasts).

Among excitatory-neuron subclasses, L2/3 intratelencephalic (IT) neurons accounted for 58.5% of the total contribution to the shared AD displacement, followed by L3/5 IT 2 and L3/5 IT 3 neurons (13.2% and 11.7%, respectively; Fig. 5b). These contributions reflected both subclass abundance and displacement magnitude. All nine subclasses showed positive projections onto the pooled AD shared direction, whereas the corresponding healthy projections were generally close to zero. Decoded shared transcriptional signatures were also highly correlated across subclasses (r = 0.69-0.97 relative to the pooled excitatory-neuron signature; Supplementary Fig. 6a), suggesting broadly consistent AD-associated remodeling across neuronal subclasses. Within patients, excitatory neurons are also shown to have lower dispersion in 6 out of 7 programs that we tested, in other word, the excitatory neurons are getting more homogenous (Supplementary Fig. 6b).

In COVID-19, the shared myeloid displacement showed a consistent interferon response accompanied by altered antigen presentation. Across Ren, COMBAT, and Stephenson cohorts, interferon-α/β signaling showed positive enrichment relative to the ten-year healthy-aging reference (z = 13.1, 8.2, and 13.5, respectively), whereas MHC class II-associated expression showed negative enrichment (z = -9.2, -10.3, and -5.2; Fig. 5c). The S100A8/A9/A12 and MS1 monocyte programs were also enriched, particularly in COMBAT and Stephenson cohorts. These changes agree with previous descriptions of interferon-responsive myeloid cells, reduced antigen presentation, and inflammatory monocyte states during acute COVID-19^44,45^, as well as the MS1 program associated with systemic infection^46^. Other inflammatory responses were less consistent: shared TNFα/NF-κB enrichment was positive in Ren and COMBAT but negative in Stephenson. Furthermore, the aging-aligned TNFα/NF-κB response was smaller than that expected from ten years of healthy aging across all three cohorts.

Interferon-associated variation was particularly pronounced between patients: the between-patient standard deviation of the decoded IFN-α/β program was 6.23-, 4.47-, and 5.53-fold higher than the healthy reference in Ren, COMBAT, and Stephenson, respectively (Fig. 5d and Supplementary Data 7). These increases were significant after multiple-testing correction in all three cohorts, whereas within-patient dispersion increased only modestly (1.12–1.30-fold). The between-patient IFN-α/β score was negatively correlated with days since symptom onset in Ren (Spearman ρ = -0.72, n = 66, q = 2 × 10^-11^) and COMBAT (ρ = -0.55, n = 46, q = 0.0001; Fig. 5e). The Stephenson cohort is excluded from this analysis because symptom onset timing is unavailable. These associations persisted after adjustment for disease severity, which showed no comparable association with interferon programs. Consistent with previous findings using MrVI^15^, these results indicate that between-patient interferon variation is associated with the timing of symptomatic infection.

In COMBAT and Stephenson, the within-patient component showed positive enrichment of the S100A8/A9/A12 program (z = 15.6 and 14.1, respectively) and negative enrichment of a non-classical monocyte signature (z = -13.8 and -14.4; Fig. 5c). To determine whether these changes reflected distinct monocyte populations, we compared CD14 and CD16 monocytes using within-subtype cell matching. CD16 monocyte fractions decreased by 12.1 percentage points in COMBAT and 10.9 percentage points in Stephenson (Fig. 5f), consistent with previously reported depletion of non-classical monocytes during COVID-19^15,44^.

Despite their reduced abundance, CD16 monocytes exhibited greater cell-level transcriptional remodeling than CD14 monocytes. Within-patient residual energy per cell increased by approximately 4 and 5 units in COMBAT and Stephenson CD16 monocytes, compared with approximately 1 unit in CD14 monocytes in both cohorts (Fig. 5f and Supplementary Data 8). The decoded CD16-identity score also decreased in CD16 monocytes (-0.09 and -0.19, respectively), accompanied by increased within-patient dispersion of calprotectin (1.46- and 1.40-fold) and CD16-identity programs (1.52- and 1.40-fold). In contrast, the corresponding dispersion ratios in CD14 monocytes were below one. All four direct CD16-versus-CD14 contrasts were significant after multiple-testing correction in COMBAT and Stephenson, but not in Ren. Together, these findings suggest that COVID-19-associated depletion of CD16 monocytes is accompanied by increased transcriptional heterogeneity and altered identity-associated expression among the remaining cells.

## Discussion

Here, we show that age-dependent changes in cross-sectional single-cell cohorts can be represented as a generative population trajectory rather than solely as a prediction of chronological age. scCody learns transport between cellular populations sampled from different individuals while using endpoint consistency to encourage sampled source populations toward a coherent target-age distribution. This formulation recovered transcriptional changes at held-out ages, enabled unseen donors to be positioned along a generated reference of healthy aging, and distinguished changes aligned with the direction of aging from those that differed from it. Across AD and COVID-19, these components differed substantially, indicating that an older predicted transcriptional age and displacement along healthy aging are not equivalent descriptions of cellular change.

Learning an aging trajectory from cross-sectional cohorts requires distinguishing relatively subtle age-dependent changes from the much larger variation shared across ages and individuals. Because cellular populations at different ages share much of their transcriptional structure, a model may achieve good population matching while generating little age-dependent change. Mean-expression profiles were reconstructed accurately by several generative models, whereas recovery of age-associated differential-expression effects more clearly separated the methods. scCody showed its strongest advantage in reconstructing the magnitude and direction of these age-dependent effects. The resulting trajectories also retained sufficient age information to localize unseen healthy donors. Meanwhile, directly supervised aging clocks remained more accurate for chronological-age prediction than trajectory localization with scCody. Age recovery provides evidence that the learned population dynamics contain biologically ordered aging information, but maximizing chronological-age prediction is not itself the objective of the generative reference. This generative reference also reproduced lineage-dependent changes in senescence and inflammatory programs. Although comparisons with prior aging studies are complicated by differences in what is measured, the most common established effects come from few group contrasts between age windows. Some other analysis relies on tracking single gene individual markers or counts of age-associated genes, or a composite program scores of genes set across age. Our program trajectories are directly comparable with the first and, to a lesser extent, the last of these. Beyond two-group contrasts, they provide continuous trajectories of program activity across adulthood. As with composite scores traced directly in observed cohorts, however, their shape cannot be reliably interpreted where donor support is thin.

Single-cell aging clocks have shown that disease, infection, and intervention can shift predicted transcriptomic age^9,27^, but the magnitude of these shifts does not establish how much of the underlying transcriptional change is aligned with healthy aging. A disease-associated state may amplify transcriptional changes that normally occur with age, acquire distinct changes that nevertheless project onto age-associated features, or combine both processes. AD contained substantial aging-aligned components together with off-trajectory remodeling, whereas COVID-19 showed comparatively weak aging alignment with more pronounced orthogonal changes. Thus, disease-associated age acceleration may capture changes shared with healthy aging without implying that the disease state as a whole follows healthy aging.

In AD patients’ excitatory neurons, both the aging-aligned and shared orthogonal displacements showed concordant enrichment with pathology-associated gene sets from the independent ROSMAP cohort^43^, linking both components to transcriptional changes associated with AD pathology. The shared displacement was broadly concordant across excitatory-neuron subclasses, with L2/3 IT neurons making the largest contribution to the population-level shift. Previous studies have also identified L2/3 IT neurons as particularly vulnerable to AD pathology^47,48^ . The reduced within-patient neuronal variation observed here may also be related to selective vulnerability^41^, although convergence toward shared disease-associated states may also be attributed. Previous single-cell studies have also identified molecular signatures shared between healthy aging and disease, including cancer.^49^

In COVID-19, previous single-cell immune aging clocks reported accelerated transcriptomic aging in monocytes,^9^ whereas our analysis revealed relatively weak aging alignment and more orthogonal components. Within these quantities, patient-level interferon variation was associated with time since symptom onset, consistent with the time-dependent interferon response reported during SARS-CoV-2 infection^50^ and myeloid patient stratification by symptom duration^15^. In COMBAT and Stephenson cohorts, CD16 monocytes showed reduced identity-associated expression and increased cell-to-cell dispersion of both calprotectin and CD16-identity programs. These observations extend previous reports of non-classical monocyte depletion^44,45^, altered inflammatory and migration-associated transcriptional programs in CD16+ monocytes^51^, and reduced inferred transitions toward non-classical monocytes^52^. The concentration of within-patient remodeling in CD16 monocytes suggests that COVID-19 also affects the distribution of transcriptional states within this population, beyond previously described changes in monocyte abundance.

Still, several improvements could be proposed for scCody. Age localization remains less accurate than chronological-age prediction by directly supervised aging clocks, and future work could improve age inference while retaining its connection to the learned healthy-aging trajectory. scCody also requires sufficient donors and cells to estimate population distributions reliably. Like other population-level optimal transport methods, these estimates are affected by finite sampling, even with entropic regularization^53,54^. This may make smaller cohorts, rare cell populations, or very fine stratification more difficult to model.

Despite these limitations, the ability to distinguish aging-aligned from off-trajectory changes could also be valuable for evaluating interventions targeting aging. Molecular aging clocks are increasingly used to assess candidate geroprotective treatments^55,56^, but a reduction in predicted molecular age does not necessarily imply reversal of the transcriptional changes associated with healthy aging. A generative healthy-aging reference could distinguish interventions that oppose age-associated transcriptional changes from those that primarily reduce off-trajectory dysregulation. These effects may produce similar changes in predicted age while reflecting different molecular responses to treatment.

## Methods

### Datasets and study design

We analyzed five single-cell or single-nucleus transcriptomic cohorts spanning healthy aging, Alzheimer’s disease (AD), and COVID-19. The Asian Immune Diversity Atlas (AIDA)^2^ was used to learn and evaluate healthy immune-aging trajectories. The PsychAD cohort (MSSM subset, which contains the most exact age donor) was used for analyses of AD^37^, and the Ren^38^, Stephenson^39^, and COMBAT^40^ cohorts were used for analyses of COVID-19. Dataset characteristics, including tissue, condition, donor and cell numbers, age coverage, and role in the study, are summarized in Supplementary Table 1.

### Healthy immune-aging cohort

The AIDA dataset comprised 625 healthy donors and approximately 1.27 million peripheral blood mononuclear cells (PBMCs). We used this cohort to train scCody, select model hyperparameters, and evaluate reconstruction and age inference in held-out donors.

For the external evaluation, we reserved all donors aged 21, 31, 41, 51, 61, and 68 years. All cells from these donors were set aside before feature selection, model training, and hyperparameter selection. The remaining 554 donors were used for model development. Age 68 was selected as the oldest held-out age because the age-71 and above group contained fewer than three donors. The external held-out populations were used only for the evaluations reported in Figs. 2 and 3. We selected 2,000 highly variable genes using Scanpy^57^ v1.11.4 with “seurat_v3”^58^ setting for model training and downstream benchmarking.

### Model Architecture

#### Gaussian-mixture latent representation

scCody represents single-cell gene-expression profiles using a Gaussian-mixture variational autoencoder (GMVAE), which aims to capture the heterogeneity across samples^15,59,60^. Let x_i_ denote the log-normalized expression profile of cell i across G = 2,000 genes. The encoder maps each cell to the mean and log-variance of a 50-dimensional Gaussian latent representation:

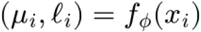

A latent state is sampled using the Gaussian reparameterization^61,62^:

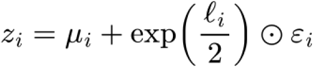

where ε_i_ is sampled from a standard multivariate normal distribution, N(0, I). The multiplication is applied element-wise. A small numerical stabilizer is included in the implementation.

The latent representation is regularized toward a learnable mixture of Gaussian components:

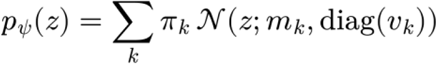

where π_k_, m_k_ and v_k_ denote the weight, mean and variance of Gaussian component k. The mixture parameters are learned jointly with the encoder and decoder.

The decoder maps the latent representation back to normalized gene-expression space:

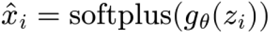

Expression reconstruction is optimized using mean-squared error across cells and genes:

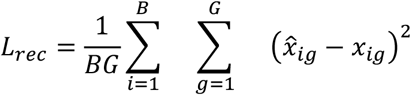

where *B* is the batch size, and *G* is the number of genes.

The latent regularization term is estimated as:

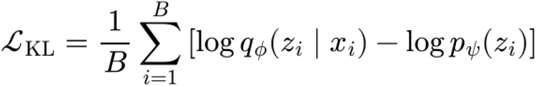

The GMVAE is trained using the combined objective:

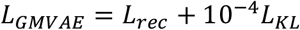

The encoder, decoder, and Gaussian-mixture prior are optimized jointly. Network architecture and the number of Gaussian-mixture components are treated as hyperparameters during model selection.

For downstream dynamics modeling, scCody uses the posterior mean as the deterministic latent state of each cell:

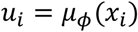

The trained GMVAE is then fixed during FlowMap training.

#### Latent dynamics model

scCody models age-associated changes in the fixed GMVAE latent space using an age-conditioned FlowMap. Chronological age was rescaled as τ = age/100. For a latent cell state u observed at source age s and target age t, the global model defines the transported state as

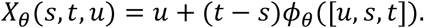

Here, φθ is a multilayer perceptron that predicts the age-dependent change in latent state. Its final layer is initialized to zero, so training begins from the identity mapping.

For scCody-CT, the transformation combines a shared mapping with a cell-type-specific residual component,

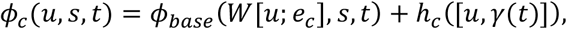

where [*u*; *e*_*c*_] denotes concatenation of the latent state and learned cell-type embedding, *W* is a learned linear projection, and *γ*(*t*) is a sinusoidal embedding^63^ of the target age. The residual heads are initialized to zero so that cell-type-specific deviations are learned from the shared mapping. Exact architectural settings are reported in Supplementary Data 9.

Each training iteration was constructed within a single sex stratum, with female and male strata sampled with equal probability. Twenty-four donors were sampled from the selected stratum and ordered by age, and cell batches were sampled from each donor to obtain latent populations U_i_, corresponding expression profiles X_i_, and donor ages τ_i_. For each target donor j, source donors were sampled bidirectionally across age. Up to three younger and three older sources were selected by dividing candidates on each side into three ranges according to absolute age difference and sampling one donor from each range. When one side contained fewer candidates, additional sources were drawn from the other side. Donors of the same age were excluded from pairing.

For source donor i and target donor j, the generated population is

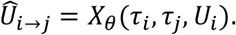

Generated and observed populations were compared using the debiased Sinkhorn divergence S,^26^ implemented with GeomLoss. The training objective combines gene-expression population matching, latent-space population matching, and endpoint consistency:

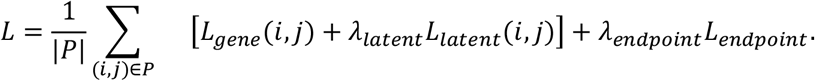

Gene-expression population matching compares decoded transported cells with the observed expression distribution of the target donor,

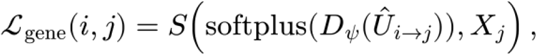

where D_ψ_ is the fixed GMVAE decoder. This loss was evaluated on decoded log1p(CP10k) expression using the squared-distance/2 transport cost. Latent-space population matching compares transported and observed target populations directly,

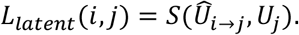

Endpoint consistency encourages different source populations transported to the same target age to produce similar endpoint distributions. For source donors a and b and common target j,

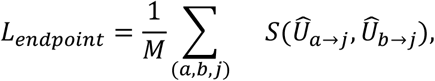

with M = 8 endpoint triplets sampled per training iteration. Source donors were sampled across the available age span and could lie on either side of the target age. Gradients were propagated through both transported endpoints. In scCody-CT, each training batch contained a single cell type, so population matching and endpoint consistency were evaluated within that cell type.

FlowMap parameters were optimized with Adam^64^ and gradient clipping. The GMVAE representation remained fixed throughout FlowMap training. Training was performed for 30 epochs with 100 iterations per epoch, with a checkpoint saved after each epoch. Sinkhorn settings, optimizer settings, architectural parameters, and tuned loss scales are reported in Supplementary Data 9.

#### Hyperparameter and checkpoint selection

Model hyperparameters were selected using three internal age-holdout folds. Each fold contained a distinct set of held-out ages across folds. For each model, 50 Optuna^65^ trials were evaluated across the three folds. Hyperparameter configurations were ranked according to the Sinkhorn divergence between generated and observed populations at the internal held-out ages, aggregated across folds. Parameters included in the search and the final values are listed in Supplementary Data 9.

After hyperparameter selection, the selected configuration was used for the final model fits. Checkpoints were saved after each training epoch, and the epoch for each fit was selected using age inference on its internal holdout donors. The checkpoint with the lowest mean absolute error was selected. Fold-specific holdout ages, selected hyperparameters, and selected epochs are reported in Supplementary Table 3 and Supplementary Data 9.

### Evaluation of held-out-age population reconstruction and age-associated expression changes

We evaluated model predictions at six held-out ages. For each held-out age, sex-matched training populations from predefined source ages spanning younger and older donors were transported to the target age and decoded to gene-expression space. 2,000 cells were generated per target age for each fitted model.

For each target age, the observed cells from held-out donors were used as the real target population. AIDA cells from the remaining training ages were used as reference populations, with donors younger than the target defining the younger reference and donors older than the target defining the older reference. Generated and observed target populations were compared separately with each reference. Analyses were performed across the full population and within B, myeloid, natural killer (NK), and T cells. With magnitude calculation of each gene calculated from Pseudobulk DE with edgeR^66^.

For each gene *g*, we calculated the observed age-associated expression change,

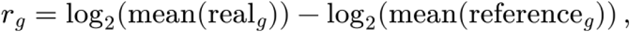

and the corresponding model-predicted change,

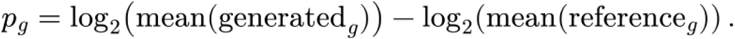

The same reference population was used for the observed and predicted contrasts. Agreement between *p*_*g*_ and *r*_*g*_ was quantified across the 2,000 modeled genes using Pearson correlation. Recovery of the greatest age-associated changes was evaluated by comparing the genes with the largest absolute observed and predicted changes. For the analyses shown in Fig. 2, overlap was calculated for the top 5% and top 10% of genes. Directional concordance was calculated as the fraction of evaluated genes for which *p*_*g*_ and *r*_*g*_ had the same sign.

Population-level reconstruction was evaluated separately by calculating the Pearson correlation between generated and observed mean expression across genes. This metric measures overall similarity between the generated and observed populations and was included as a separate component of the benchmark summary.

To estimate the variability of age-associated expression effects in the observed data, held-out donors at each target age were randomly divided into two groups. Expression changes relative to the same age-specific reference population were calculated independently in each half and compared using the same evaluation metrics. This procedure was repeated 500 times. The resulting distribution provided the split-half reference shown in Fig. 2. The same procedure was applied within each of the four major immune cell types.

For models that did not directly retain AIDA-compatible cell-type labels in their generated populations, labels were assigned using a nearest-neighbor classifier trained on 50,000 reference AIDA cells represented by 50 principal components with 15 nearest neighbors. Cell-type-resolved expression changes were then calculated using reference, observed, and generated cells from the same cell type.

Results were summarized across the six held-out ages and three fitted models. Global metrics are reported as the mean and standard deviation across age–fit combinations. For cell-type macro summaries, B, myeloid, NK, and T cells were weighted equally, and younger- and older-reference comparisons were likewise given equal weight.

### Trajectory-based age inference

We inferred donor age by identifying the position along the learned aging trajectory that best matched the donor’s observed cell population. For each model fit, cells from a held-out donor were represented by their GMVAE posterior means. Five fixed source ages spanning the training range were used, with source populations formed by pooling training cells of the same sex as the target donor. Up to 2,000 cells were sampled from each source and target population. Candidate ages were evaluated at one-year intervals across the training age range, permitting transport toward both younger and older ages.

For source population X_s_ observed at age a_s_, cells were transported to each candidate age a and compared with the target population Y:

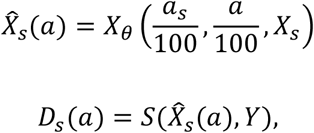

where S is the latent-space Sinkhorn divergence defined above. The same sampled source cells were used for all candidate ages, and source and target populations were matched in size for each comparison. The estimate from source s was the age minimizing this distance,

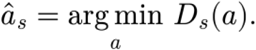

Age estimates were averaged across the five source populations within each fit and subsequently across the three model fits to obtain the final donor-level estimate.

Cell-type-specific age inference used the same procedure after restricting the source and target populations to the corresponding lineage. scCody-G used the global transport map, whereas scCody-CT used the corresponding cell-type-conditioned map. The source ages used for each fit are provided in Supplementary Table 3.

Age-inference performance was evaluated using mean absolute error and Pearson correlation between inferred and chronological age across the held-out donors. Confidence intervals were obtained by bootstrap resampling donors 2,000 times.

### Supervised age-prediction models

We compared trajectory-based age inference with supervised Lasso, ElasticNet and tAge models trained on donor-level pseudobulk expression. All models used the same three development splits and 2,000 genes described above. Raw counts were summed into metacells, either across all cells from each donor or within donor–cell-type combinations. Metacells containing fewer than five cells were excluded, and genes were retained if detected in at least 20% of the training metacells.

Normalization parameters were estimated from the training metacells of each fit and applied unchanged to held-out data. DESeq relative log expression (RLE) size factors^67^ were calculated from gene-wise geometric means, after which normalized expression was computed as

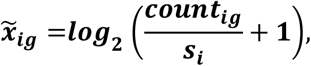

where s_i_ is the RLE size factor for metacell^9,28,68,69^ i. Expression values were then standardized using training-set gene means and standard deviations.

Following the tAge strategy, expression and chronological age were centered relative to a young reference population. For each modeled stratum, the reference consisted of training metacells from donors aged 30 years or younger. The reference expression vector m_k_ was defined as the gene-wise median across these metacells, and the corresponding reference age a_k_ as their median chronological age. Models were trained to predict relative age from relative expression,

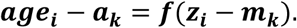

For cell types with fewer than five young-reference metacells, the reference expression was estimated from all eligible young metacells. Predicted chronological age was obtained by adding a_k_ to the model output.

All supervised models used linear ElasticNet regression, with the Lasso corresponding to the pure L1 case. Hyperparameters were selected using the same three internal age-holdout folds used for model development. Candidate α and L1-ratio combinations were fitted separately within each fold and ranked by mean donor-level MAE across the three internal holdouts. The selected hyperparameters were subsequently refitted within each of the three development splits, and predictions were averaged across fits. Search ranges and selected values are reported in Supplementary Data 9.

The supervised models differed in how metacells were defined and how regression models were shared across cell types. Global Lasso and ElasticNet used one metacell per donor and fitted a single model. The tAge model used donor–cell-type metacells but fitted one shared regression model across cell types, with each cell type centered on its own young reference. Its donor-level age estimate was the cell-count-weighted average of the available lineage predictions. Cell-type-specific ElasticNet models instead fitted an independent regression for each lineage, with hyperparameters selected separately by lineage. Cell-type-specific Lasso models followed the same procedure with the L1 ratio fixed at 1.

Performance was evaluated using MAE, Pearson and Spearman correlation, R^2^, and calibration slope between predicted and chronological age.

### Immune-aging pathway analysis

To examine whether scCody recapitulates established immune-aging programs, we projected observed cells across the learned aging trajectory using the trained age-conditioned FlowMap. For a cell with expression profile *x* observed at source age *a*_*s*_, its latent representation was obtained using the frozen GMVAE encoder and transported directly to a target age *a*_*t*_, followed by decoding into gene-expression space:

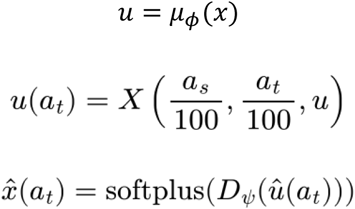

For each target age, the transformation was applied directly to the original source population rather than sequentially propagating cells through intermediate ages. The same source cells were used across target ages, and decoded expression was averaged within each donor–cell-type group to obtain donor-level expression profiles. Analyses were performed within T, myeloid, B, NK, dendritic, and platelet populations.

For the immune-aging pathway analysis presented here, we used held-out 31-year-old AIDA donors as the source population and projected their cells across the aging trajectory at one-year increments.

We evaluated six predefined gene programs primarily from MSigDB^70^ and Reactome^71^: Hallmark Inflammatory Response, Hallmark TNFα signaling via NFκB, Reactome Cellular Senescence, Reactome Senescence-Associated Secretory Phenotype, Reactome Signaling by the BCR, and a curated TCR signaling program. BCR and TCR programs were evaluated in B and T cells, respectively, whereas the remaining programs were evaluated across all analyzed lineages. Only genes represented among the 2,000 modeled genes were included. Program details are available in Supplementary data 3.

Program activity was calculated using an expression-matched control-gene procedure. For each cell type and gene program, the 2,000 genes were ranked by mean expression across real AIDA donor pseudobulks and divided into expression bins.^72^ Control genes were sampled from the same bins as the program genes using a fixed random seed, and the resulting control set was held constant across real and generated data and across model fits. For an expression profile E, program activity was defined as

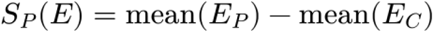

where P denotes the program genes and C the corresponding expression-matched control genes. Scores were calculated from donor-level mean log1p(CP10k) expression profiles.

For each projected donor–cell-type group g, changes in program activity were expressed relative to its reconstructed starting baseline:

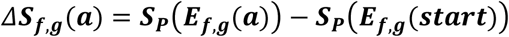

Projected changes were averaged first across donors within each fit and then across the three model fits. For uncertainty bands, each donor’s projected change was first averaged across fits, and 95% confidence intervals at each age were calculated as the donor-level mean ± 1.96 × s.e.m. The same gene-program scoring framework was applied to real AIDA donor pseudobulks to obtain the corresponding cross-sectional age-associated changes.

### Disease-associated age shifts

Disease-associated age shifts were evaluated in an AD brain cohort and three COVID-19 PBMC cohorts. For AD, a scCody model was trained using healthy D11 donors and applied to 212 AD donors and healthy holdout donors with available projections. For COVID-19, the three fitted AIDA models were applied to the Ren, COMBAT, and Stephenson cohorts. Each COVID-19 dataset was restricted to the AIDA 2,000-gene panel, with genes absent from a cohort assigned the corresponding healthy-reference mean.

Donor age was inferred using the trajectory-localization procedure described above. Briefly, each donor was localized independently from five age-spanning source populations by identifying the candidate age with minimum latent-space Sinkhorn divergence, and source-specific estimates were averaged. COVID-19 estimates were additionally averaged across the three AIDA model fits. Global and cell-type-specific ages were calculated using the corresponding whole-population or lineage-restricted cells.

For donor i, the raw age shift was defined as

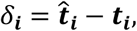

where 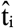 is the inferred age, and t_i_ is the chronological age. To account for age-dependent bias in model predictions, each disease donor was compared with healthy donors of similar chronological age. The main analysis matched controls within ±5 years of the disease donor; COVID-19 controls were drawn from the same cohort. AD analyses required at least three matched controls and COVID-19 analyses at least one. The disease-associated effect for donor i was

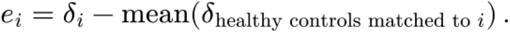

Controls could contribute to more than one disease donor. Cohort-level effects were calculated as the mean of e_i_ across disease donors. Confidence intervals were obtained by bootstrap resampling disease donors, and significance was assessed using a one-sample t-test against zero with Benjamini–Hochberg correction within each cohort and analysis scope.

Supervised age estimates were evaluated using the ElasticNet framework described above. For COVID-19, the frozen AIDA-trained supervised models were applied to each cohort and combined across the three fits using the same procedure as for healthy AIDA donors. For AD, a separate ElasticNet model was trained on healthy D11 donors using donor-by-cell-class pseudobulk profiles. Counts were normalized to CP10k, log1p-transformed, and standardized before ElasticNet regression. Disease effects were calculated using the same local healthy-matching procedure as for scCody.

Agreement between scCody and ElasticNet was quantified using Lin’s concordance correlation coefficient (CCC)^73^ on each donor’s raw age shift from the two methods. For paired estimates x and y,

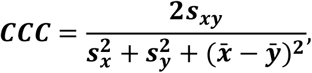

where sx^2^ and sy^2^ are sample variances, and s_xy_ is the sample covariance. CCC was calculated separately by cohort, analysis scope, and cell type using donors scored by both methods.

### Construction of disease-associated and healthy-aging displacement fields

To determine how disease-associated remodeling relates to healthy aging, we constructed disease-associated and predicted aging displacement fields in the scCody latent space. Each disease donor was paired with healthy reference donors from the same cohort and sex whose chronological ages differed by at most five years. Cell-type analyses were restricted to the same lineage in both donors. For each donor pair, equal numbers of healthy and disease cells were sampled and matched by exact optimal transport, minimizing the total squared Euclidean distance between their latent states. For healthy reference cell *x*_*i*_ and its matched disease cell *yπ*(*i*), the disease-associated displacement was

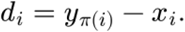

For the same reference cells, scCody predicted the corresponding aging displacement based on the disease donor’s inferred age shift relative to its matched healthy controls,

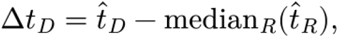

where 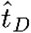 and 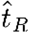 denote trajectory-derived age estimates. The predicted aging displacement for each reference cell was then

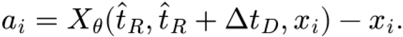

Both displacement fields originated from the same healthy reference cells, allowing disease-associated remodeling to be compared with the predicted aging movement corresponding to the donor’s inferred age shift. Pairs whose projected target ages fell outside the fitted age range were excluded.

A healthy-aging reference was constructed from same-sex healthy donor pairs within each cohort. The primary H10 reference paired younger and older donors separated by 8–12 years. For each healthy pair, the observed displacement was obtained by optimal-transport matching from younger to older donor cells, while the corresponding aging displacement was predicted between their chronological ages,

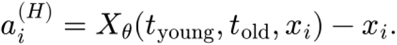

For AD, H10 target donors were restricted to the 74-donor healthy holdout excluded from brain-model training. For COVID-19, healthy comparisons used controls from the corresponding cohorts, independent of AIDA model training. The same alignment and residual-decomposition analyses were applied to disease and healthy pairs.

### Alignment of disease-associated displacement with healthy aging

For each matched donor pair *d*_*pi*_, we compared the disease displacement with the predicted aging displacement *a*_*pi*_ of the same healthy reference cell . These cell-specific aging displacements collectively define the healthy-aging field. We calculated

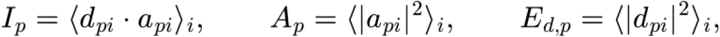

where ⟨·⟩_*i*_ denotes the mean across matched cells. A single least-squares coefficient was estimated for the entire donor pair,

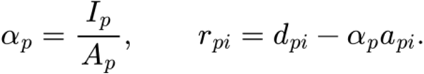

Thus, *α*_*p*_*α*_*pi*_ is the component captured by a common rescaling of the cell-specific aging field, and *r*_*pi*_ is the remaining displacement. Orthogonality is defined across the population as, ⟨*r*_*pi*_ ⋅,*a*_*pi*_ ⟩_*i*_*=*0. We quantified the signed magnitude of aging alignment as

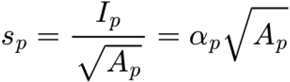

The signed aligned RMS, (s_p), is the signed RMS length of the aging-aligned component and is termed **aging-aligned displacement** in the Results. Positive and negative values indicate alignment with and opposition to the model-predicted aging movement, respectively.

These normalized projection quantities were evaluated for ***A***_*p*_ >10^−12^. For numerically negligible aging fields, we set *alpha*_*p*_ = 0, retained the full disease displacement as the residual, and recorded the signed aligned RMS as undefined.

The aligned and orthogonal energies were

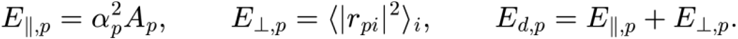

where the signed aligned RMS is defined as 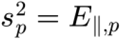. Energies are then partitions of the magnitude of cellular remodeling, while the signed aligned RMS also retains its direction relative to the aging field.

### Donor-level summaries and comparison with healthy aging

For the displacement comparison in Fig. 4d, signed aligned RMS was calculated for each donor pair before aggregation. Pair scores were averaged over matched healthy references within each target donor and then equally across target donors. Residual energies were averaged using the same weighting before conversion to RMS. COVID-19 summaries gave each cohort equal weight.

The mean signed score describes net aging alignment. Its square is distinct from the mean aligned energy, because positive and negative pair scores can offset one another during averaging.

Relative aging alignment and excess orthogonal displacement were reported as

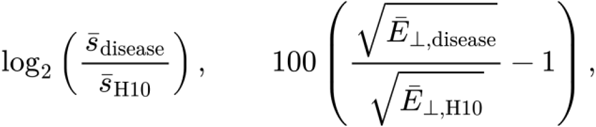

respectively. The first compares signed aging-aligned displacement with the healthy-aging reference; the second compares orthogonal displacement on an RMS scale. The nonpositive global COVID-19 signed score was represented by a boundary marker annotated with its untransformed value.

### Shared, between-patient and within-patient remodeling

We further partitioned residual energy according to its organization across patients and cells. For target donor (j), we calculated the mean residual vector and variation around that mean,

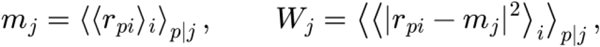

where ⟨⋅⟩ denotes an equal-weight average over the reference-donor comparisons for donor (j). With 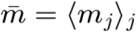, equal weighting of target donors gives

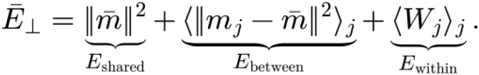

The shared component measures the cohort-average residual shift, the between-patient component measures differences between patient means, and the within-patient component measures the remaining variation within each patient. The latter includes variation across cells and across matched healthy-reference comparisons. These components are termed **shared shift, between-patient variation**, and **within-patient variation** in the Results.

The same decomposition was applied to the healthy-aging reference. For each component *k* (shared, between-patient, or within-patient), its contribution to the disease-associated change in total orthogonal energy was expressed as

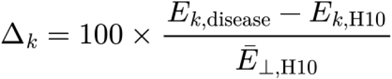

Here, *E*_*k,disease*_ and *E*_*k,H10*_ denote the corresponding component energies in disease and the healthy ten-year reference, respectively, and 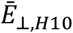 denotes the mean total orthogonal energy in the healthy reference. Each component’s contribution sums to the total percentage change in orthogonal energy relative to healthy aging.

### Functional interpretation of disease-associated displacement

To characterize the transcriptional effects of the aging-aligned and non-aging components of disease-associated displacement, we decoded a sequence of latent states for each optimal-transport-matched healthy-disease cell pair. Let *z* denote the latent state of a healthy source cell; *z*′ its matched disease cell; *αa* the aging-aligned displacement; 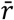 the shared non-aging residual across disease donors; and *r*_*D*_ the donor-specific mean residual. Let denote the fixed GMVAE decoder, including its output transformation. The decoded expression profiles were defined as

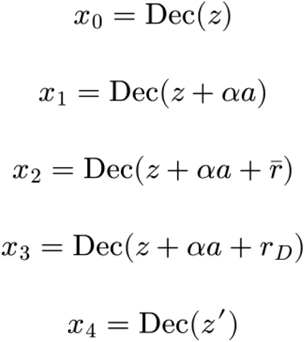

The corresponding expression-space effects were *x*_1_ − *x*_0_ (aging-aligned), *x*_2_ − *x*_1_ (shared non-aging), *x*_3_ − *x*_2_ (between-patient), and *x*_4_ − *x*_3_ (within-patient). These effects sum to the total matched expression difference, *x*_4_ − *x*_0_.

For each expression-space component, gene-program activity was calculated at the cell level as the mean effect across genes in the program minus the mean effect across all modeled genes:

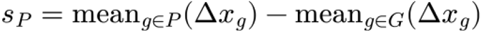

where *P* denotes the program gene set and *G* the available modeled genes. Cell-level scores were averaged within each matched donor pair, then across pairs for each donor, and subsequently across model fits. Donor-cell-type groups were required to contain at least 30 cells within a fit. Gene programs were assembled from published MSigDB sets and curated literature-derived gene lists. For AD pathology-associated gene sets, we used the DecontX/RUVr-adjusted excitatory-neuron differential-expression results for global AD pathology from Mathys et al. (2023). Genes were retained if they showed significant associations (within-subtype BH-adjusted P < 0.05) in the same direction in at least 7 of 14 excitatory-neuron subtypes, with no significant association in the opposite direction. The resulting gene sets were intersected with the modeled genes.

For the AD EN and COVID-19 myeloid program analyses, each decoded component was compared with the corresponding +10-year healthy reference described above. For each gene *g*, the difference between its mean disease and healthy donor effects was denoted Δ_*g*_. The observed program contrast, *Q*_*P*_, was calculated as the mean Δ_*g*_ across program genes minus the mean across all available genes. To account for gene-set size, this contrast was standardized against 2,000 randomly sampled gene sets of the same size:

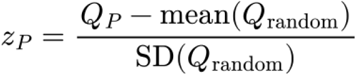

where Q denotes the program contrast. Random sets were drawn from the measured genes for COVID-19 and from the complete modeled panel for AD and were fixed for each cohort and gene-set size. Significance was assessed by 10,000 two-sided donor-label permutations, followed by Benjamini– Hochberg correction across programs within each cohort and decoded component.

To identify the cellular sources of the shared AD displacement, EN cells were grouped by the subclass of the healthy source cell. The shared residual direction was defined as

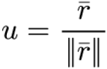

and each cell residual was projected onto this direction as ***r***_***i***_ ⋅ ***u***. Disease and healthy-reference cells were projected onto the same direction, and donor-level means were calculated for groups containing at least 10 cells. Confidence intervals were obtained from 2,000 donor bootstrap resamples. The contribution of subclass l to the shared displacement was summarized as its AD cell fraction multiplied by its donor-mean projection and expressed relative to the sum across subclasses represented by at least five AD and five healthy donors. Cell fractions and projection magnitudes were recomputed jointly within each bootstrap sample.

For COVID-19, heterogeneity of program effects was quantified separately between and within patients. Between-patient dispersion was the sample standard deviation of donor-level program scores, whereas within-patient dispersion was the mean across donors of the cell-level standard deviation, using groups with at least 30 cells. Disease-to-healthy dispersion ratios were calculated for each program and component. Confidence intervals were obtained from 10,000 donor bootstrap resamples; enrichment of dispersion in COVID-19 was tested using one-sided permutation tests followed by Benjamini– Hochberg correction within each cohort and dispersion level.

Associations between interferon-related remodeling and time since symptom onset were evaluated in COVID-19 donors with a single sampled time point. Between-patient program scores were averaged within matched pairs and across model fits to obtain one value per donor. Associations with days since symptom onset were assessed using Spearman correlation. To account for disease severity, we additionally fitted a rank-based linear model,

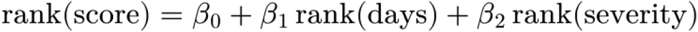

*P* values were corrected using the Benjamini-Hochberg procedure across the three interferon programs within each cohort, decoded component, and statistical test. Regression lines shown in the figure were fitted against log-transformed days for visualization.

Finally, CD14 and CD16 monocyte remodeling was examined using harmonized subtype annotations. Target-cell subtype labels were assigned after optimal-transport matching. Changes in subtype abundance were calculated from the proportions of CD14 and CD16 cells among all myeloid target cells. Analyses of residual energy and program activity included matched cells with concordant source and target subtype annotations. COVID-19-versus-healthy contrasts were evaluated for subtype abundance, within-patient residual energy, the Azimuth CD16 program score, and within-patient dispersion of GO:1990660 and the Azimuth CD16 program. Per-subtype differences were assessed by permutation testing with Benjamini-Hochberg correction. CD16-versus-CD14 contrasts were calculated as differences for residual energy and mean program effects and as log ratios for dispersion ratios. Confidence intervals and two-sided *P* values were obtained from 10,000 paired donor bootstrap resamples, with COVID-19 and healthy donors resampled separately, and both monocyte subtypes recomputed within each bootstrap sample. *P* values were corrected across the four outcomes within each cohort.

## Code availability

All software development, data processing, and statistical analyses were performed using Python (version 3.10.18) and R (version 4.4.1) along with their respective packages. The full source code for scCody is openly available at https://github.com/jaspershen-lab/sccody. Scripts used for data processing, statistical analysis, and figure generation are available at https://github.com/jaspershen-lab/sccody_manuscript.

## Data availability

All datasets analyzed in this study and used as demonstration data were obtained from previously published sources^2,37–40^, with dataset characteristics and accession identifiers provided in Supplementary Table 1. Additional source data supporting the findings of this study are provided in the Supplementary Data.

## Acknowledgments

This work was supported by the Nanyang Assistant Professorship start-up funding awarded to Xiaotao Shen by the Lee Kong Chian School of Medicine and the School of Chemistry, Chemical Engineering and Biotechnology at Nanyang Technological University, Singapore. Additional support was provided by a Ministry of Education (MOE) Academic Research Fund Tier 1 grant awarded to Xiaotao Shen (Grant No. 025402-00001). The authors acknowledge the use of ChatGPT (OpenAI) for language editing and grammatical refinement during manuscript preparation.

## Author contributions

J.K. and X.S. jointly conceived the project. X.S. and J.K. supervised the overall study. J.K. designed the methodology and developed the algorithm. J.K. analyzed all the datasets. J.K. and X.S. jointly prepared all figures and wrote the manuscript. J.C. and S.Z. reviewed and edited the manuscript. All authors reviewed and approved the final version of the manuscript.

## Additional information

Correspondence and requests for materials should be addressed to X.S.

