## Supplementary for "scCody distinguishes aging-aligned from divergent disease-associated cellular remodeling"

**scCody disentangles aging progression from off-trajectory dysregulation in cross-sectional single-cell cohorts**

**Supplementary Figures**

1. Supplementary Fig. 1. Cell-type annotation, age stratification and highly variable gene selection in the AIDA healthy reference cohort.
2. Supplementary Fig. 2. Extended benchmarking of population reconstruction and age-associated differential expression at held-out ages.
3. Supplementary Fig. 3. Extended evaluation of trajectory-based donor-age inference and ablation of endpoint-consistency and latent optimal-transport regularization.
4. Supplementary Fig. 4. Donor-age inference by the brain model in held-out healthy PsychAD donors and its agreement with ElasticNet.
5. Supplementary Fig. 5. Disease-associated age shifts in each COVID-19 cohort estimated by scCody and ElasticNet.
6. Supplementary Fig. 6. Subclass consistency and patient-level dispersion of AD-associated displacement in excitatory neurons.
7. Supplementary Fig. 7. Severity-adjusted association of interferon remodeling with symptom duration, and CD14/CD16 monocyte comparisons in the Ren cohort.

**Supplementary Tables**

1. Supplementary Table 1. Dataset Characteristics, roles and data sources of the single-cell and single-nucleus cohorts analyzed in this study.
2. Supplementary Table 2. Ablation of endpoint-consistency and latent optimal-transport regularization in scCody-G and scCody-CT.
3. Supplementary Table 3. Internal age-holdout folds, selected checkpoints and settings for trajectory-based age inference.

**Supplementary Data**

1. Supplementary Data 1. Population reconstruction and differential-expression benchmark metrics by model, held-out age, model fit and cell type.
2. Supplementary Data 2. Trajectory-derived and supervised age estimates for held-out healthy AIDA and PsychAD donors.
3. Supplementary Data 3. Gene programs used for immune-aging and disease-displacement analyses.
4. Supplementary Data 4. Disease-associated age shifts, concordance between scCody and ElasticNet in Alzheimer’s disease and COVID-19 cohorts.
5. Supplementary Data 5. Aging-aligned, orthogonal, shared, between-patient and within-patient displacement estimates relative to the ten-year healthy-aging reference.

6. Supplementary Data 6. Gene-program enrichment of decoded displacement components in AD excitatory neurons and COVID-19 myeloid cells.
7. Supplementary Data 7. Between- and within-patient dispersion of decoded programs, and associations of interferon programs with time since symptom onset in COVID-19.
8. Supplementary Data 8. CD14 and CD16 monocyte comparisons of abundance, residual energy and program dispersion across COVID-19 cohorts.
9. Supplementary Data 9. Model architectures, optimization settings and hyperparameter search spaces with selected values for scCody and comparison generative models.

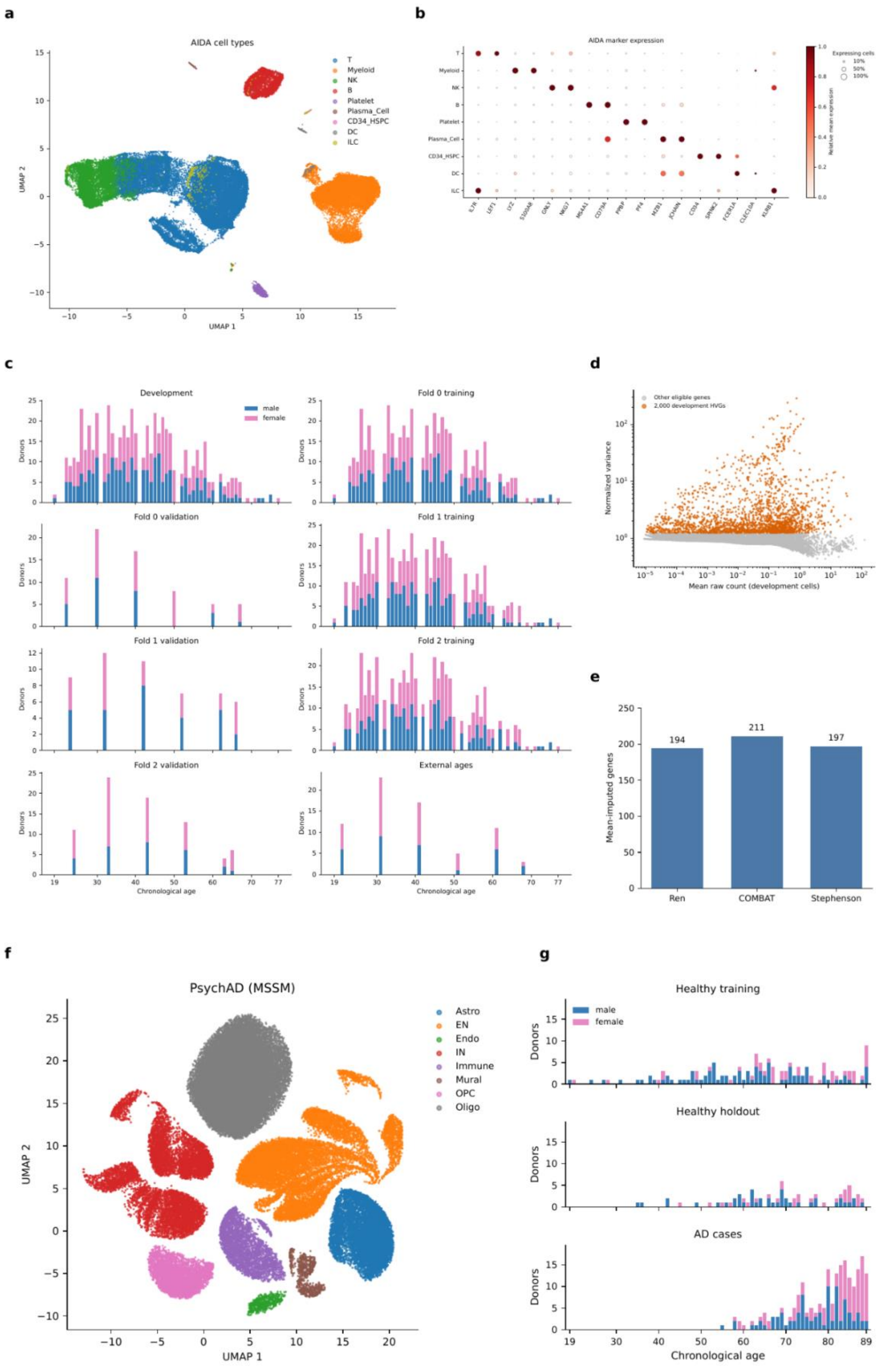
a
AIDA cell types
T
Myeloid
NK
B
Platelet
Plasma_Cell
CD34_HSPC
DC
ILC
UMAP 1
UMAP 2
b
AIDA marker expression
Expressing cells
10%
50%
100%
Relative mean expression
IL7R
LEF1
LYZ
S100A8
GNLY
NKG7
MS4A1
CD79A
PPBP
PF4
MZB1
JCHAIN
CD34
SPINK2
FCER1A
CLEC10A
KLRB1
c
Development
male
female
Donors
Fold 0 training
Fold 0 validation
Fold 1 training
Fold 1 validation
Fold 2 training
Fold 2 validation
External ages
Chronological age
d
Other eligible genes
2,000 development HVGs
Normalized variance
Mean raw count (development cells)
e
Mean-imputed genes
194
211
197
Ren
COMBAT
Stephenson
f
PsychAD (MSSM)
Astro
EN
Endo
IN
Immune
Mural
OPC
Oligo
UMAP 1
UMAP 2
g
Healthy training
male
female
Donors
Healthy holdout
AD cases
Chronological age

**Supplementary Fig. 1 | Overview of the healthy reference, COVID-19 and Alzheimer's disease (AD) cohorts.** (a) UMAP of AIDA PBMCs coloured by major cell type. (b) Expression of marker genes across AIDA cell types. Dot size indicates the fraction of expressing cells, and colour indicates mean expression scaled to the maximum across cell types. (c) Donor counts by chronological age and sex in the development set, the training and validation splits of the three internal age-holdout folds, and the external held-out ages (21, 31, 41, 51, 61 and 68 years). (d) Selection of 2,000 highly variable genes from development cells. Highly variable genes are shown in orange. (e) Number of modeled genes absent from each COVID-19 cohort and assigned the healthy-reference mean. (f) UMAP of PsychAD (MSSM) nuclei coloured by cell class. (g) donor counts by age and sex for healthy training donors, healthy holdout donors and AD cases.

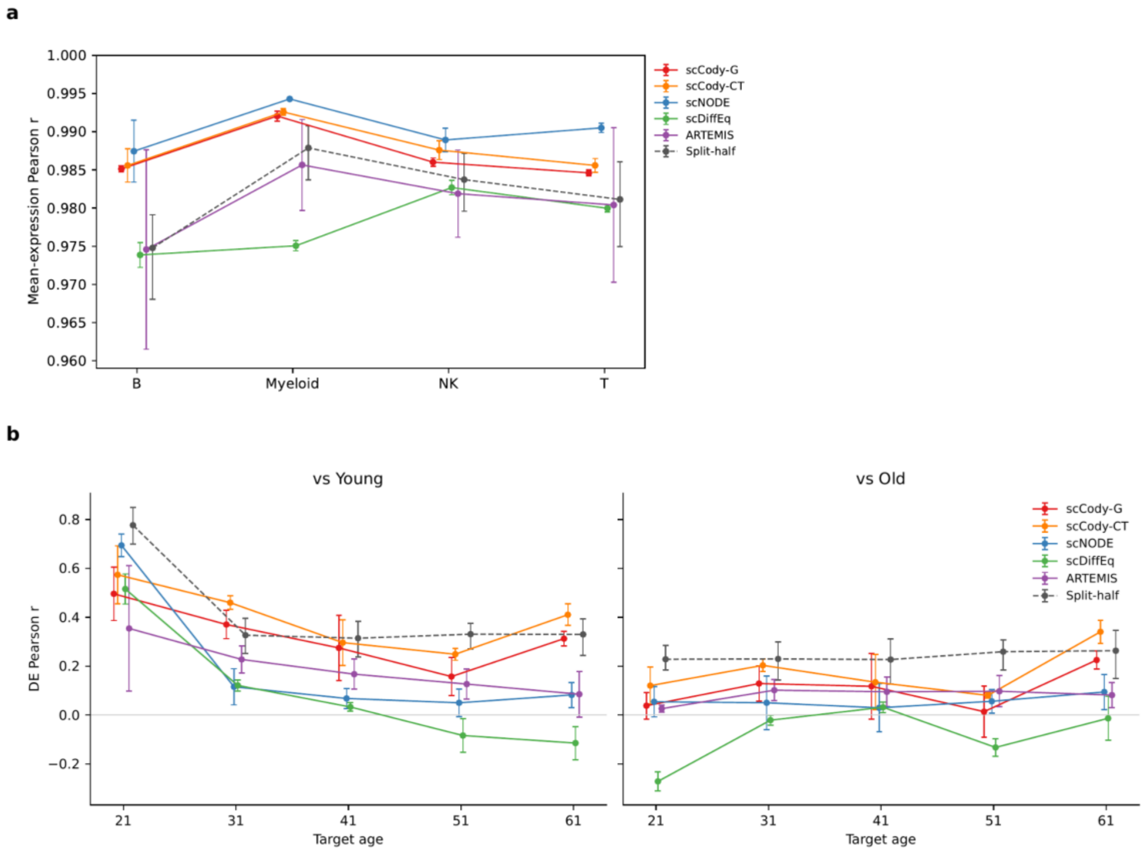


**Supplementary Fig. 2 | Population reconstruction by lineage and recovery of age-associated differential expression across held-out ages.** (a) Pearson correlation across genes between generated and observed mean expression at held-out ages within B, myeloid, NK, and T cells. Points show means across held-out ages and model fits, and bars show s.d. (b) Pearson correlation between predicted and observed gene-wise log2 fold changes at each held-out age, relative to younger (left) and older (right) reference populations. Points show means across three model fits, and bars show s.d. In both panels, split-half (grey dashed) shows agreement between two random halves of the observed held-out donors (500 repeats); points show the mean and bars show 95% confidence intervals.

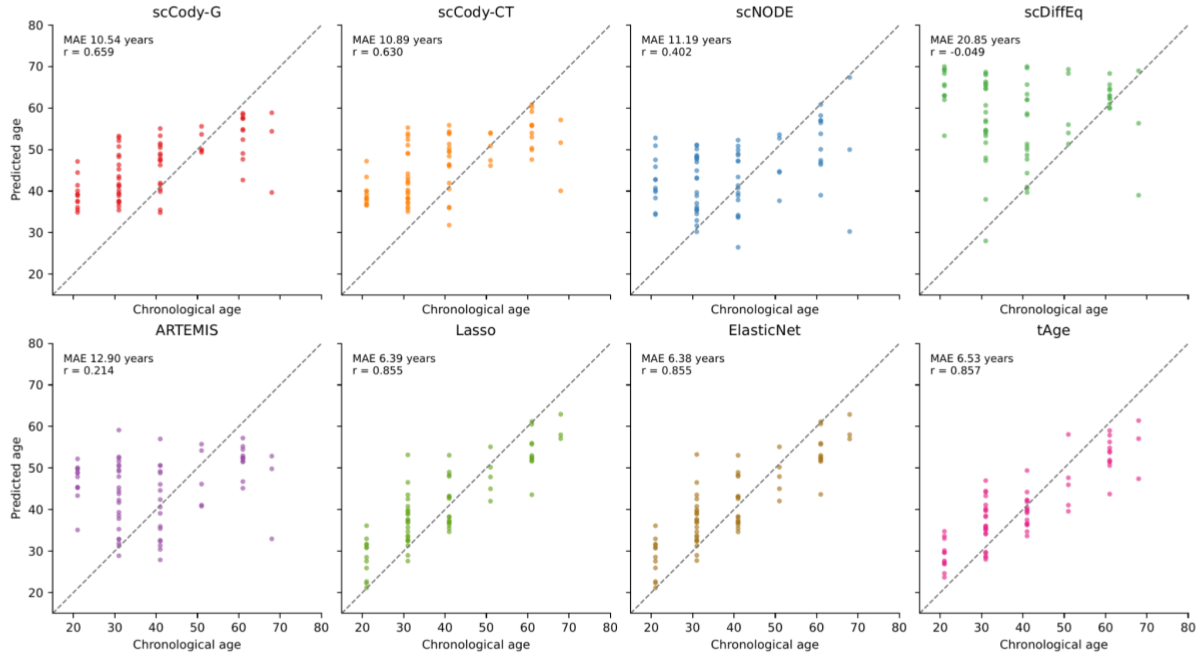


**Supplementary Fig. 3 | Donor-age inference in held-out healthy AIDA donors.** Predicted versus chronological age for the 71 external held-out donors, for scCody-G, scCody-CT, scNODE, scDiffEq, ARTEMIS, Lasso, ElasticNet and tAge. Each point is a donor. Estimates are averaged across three-fold model ensembles, and for generative models also across five source populations. Dashed lines indicate identity. MAE and Pearson r are shown in each panel.

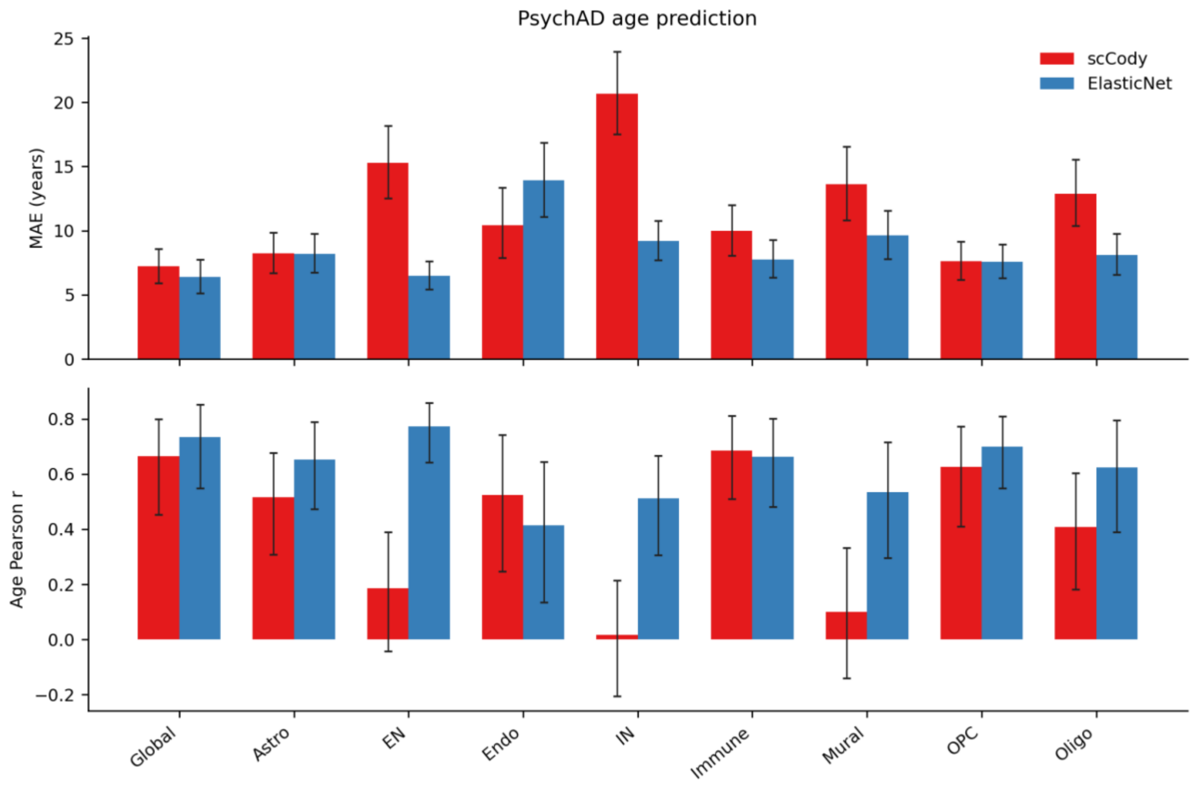


**Supplementary Fig. 4 | Donor-age inference by the brain model in held-out healthy PsychAD donors. (a,b)** MAE (**a**) and Pearson correlation between predicted and chronological age (**b**) for scCody and ElasticNet in 74 held-out healthy PsychAD donors, computed globally and within each cell class. Both models were trained on the same healthy training donors. Error bars show 95% confidence intervals.

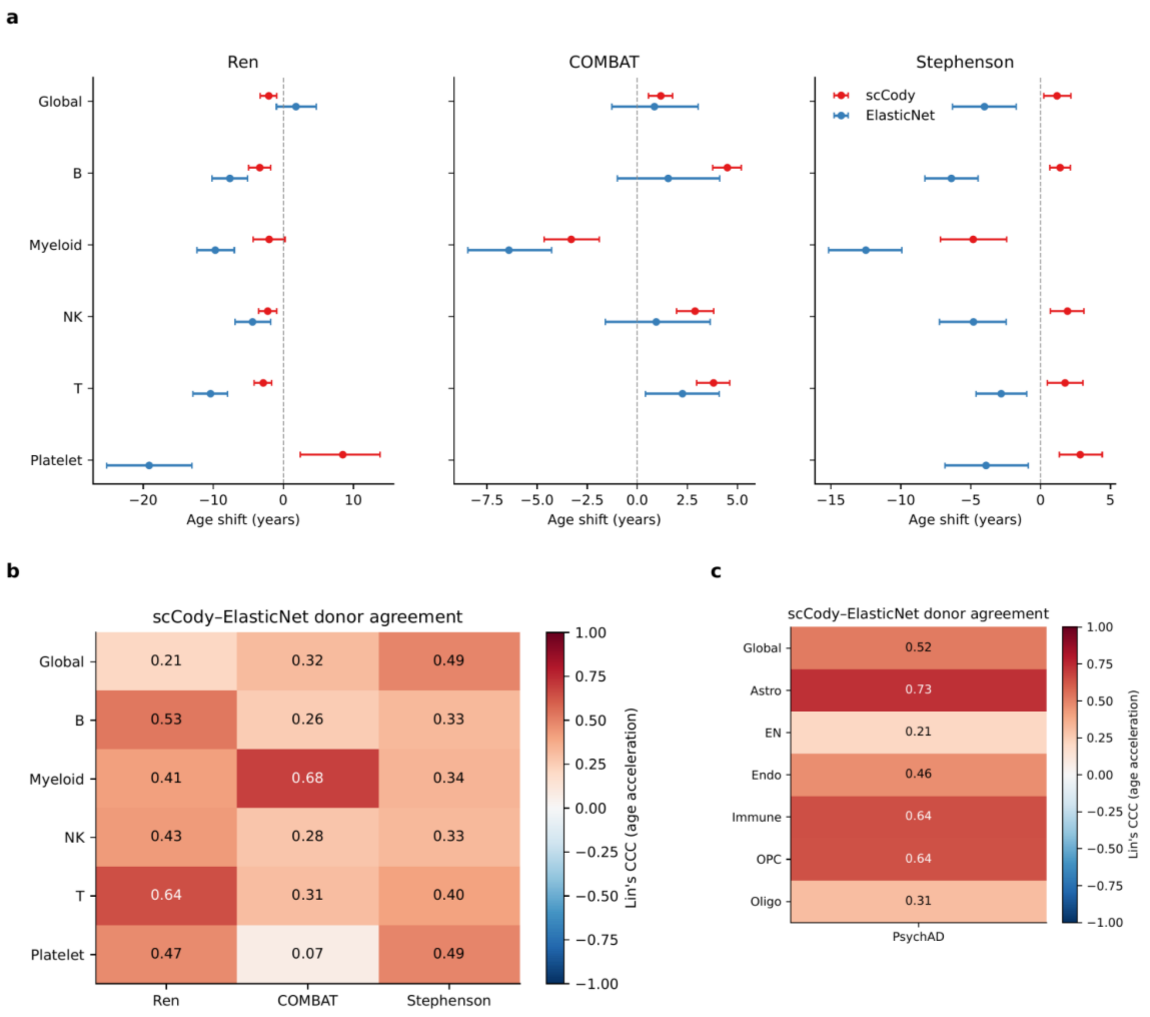


**Supplementary Fig. 5 | Disease-associated age shifts in COVID-19 cohorts and concordance between scCody and ElasticNet.** (a) Age shifts estimated by scCody (red) and ElasticNet (blue) in the Ren, COMBAT, and Stephenson cohorts, globally and within each cell type. The age shift is the gap between inferred and chronological age in patients minus the same gap in healthy donors from the same cohort within ± 5 years of age. Points show means, and bars show 95% bootstrap confidence intervals. (b,c) Lin's concordance correlation coefficient between scCody and ElasticNet donor-level age acceleration in each COVID-19 cohort (b) and in PsychAD (c), globally and within each cell type.

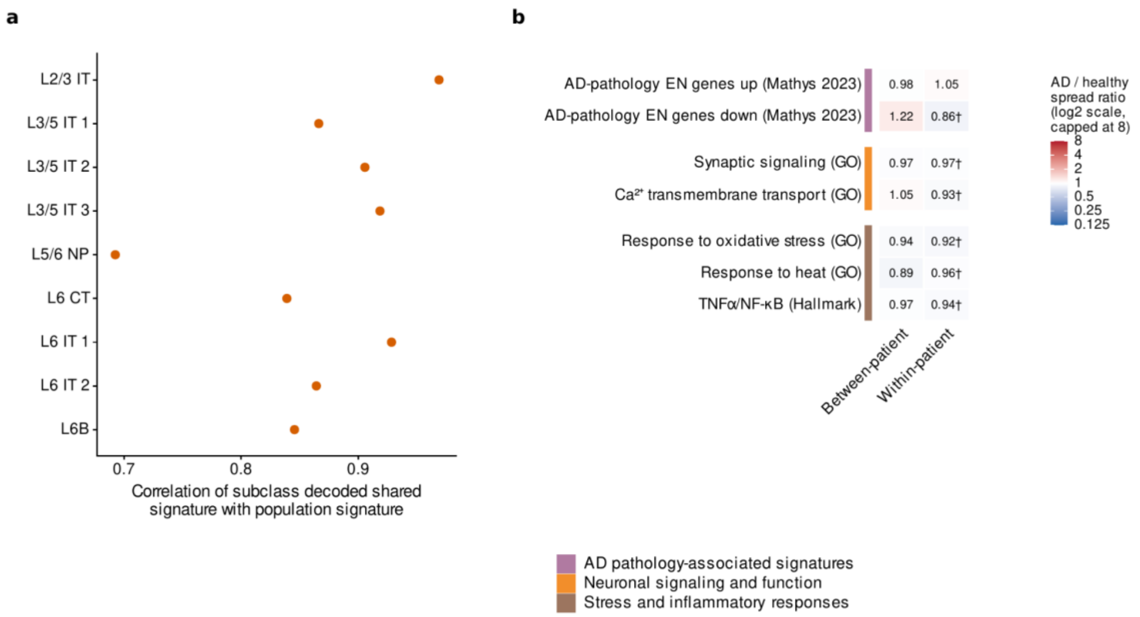


**Supplementary Fig. 6 | Subclass consistency and patient-level dispersion of AD-associated displacement in excitatory neurons.** (a) Pearson correlation between the decoded shared AD signature of each excitatory-neuron subclass and the signature of the pooled excitatory-neuron population. (b) Dispersion of decoded gene-program effects in AD relative to the ten-year healthy-aging reference, shown as AD/healthy standard-deviation ratios. Between-patient dispersion is the s.d. of donor-level program scores, and within-patient dispersion is the mean within-donor s.d. Colour shows ratios on a log2 scale, and coloured bars indicate program groups. † indicates significantly lower dispersion in AD (ratio < 1; one-sided permutation test, Benjamini–Hochberg q < 0.05), and * indicates significantly higher dispersion (ratio > 1; same test).

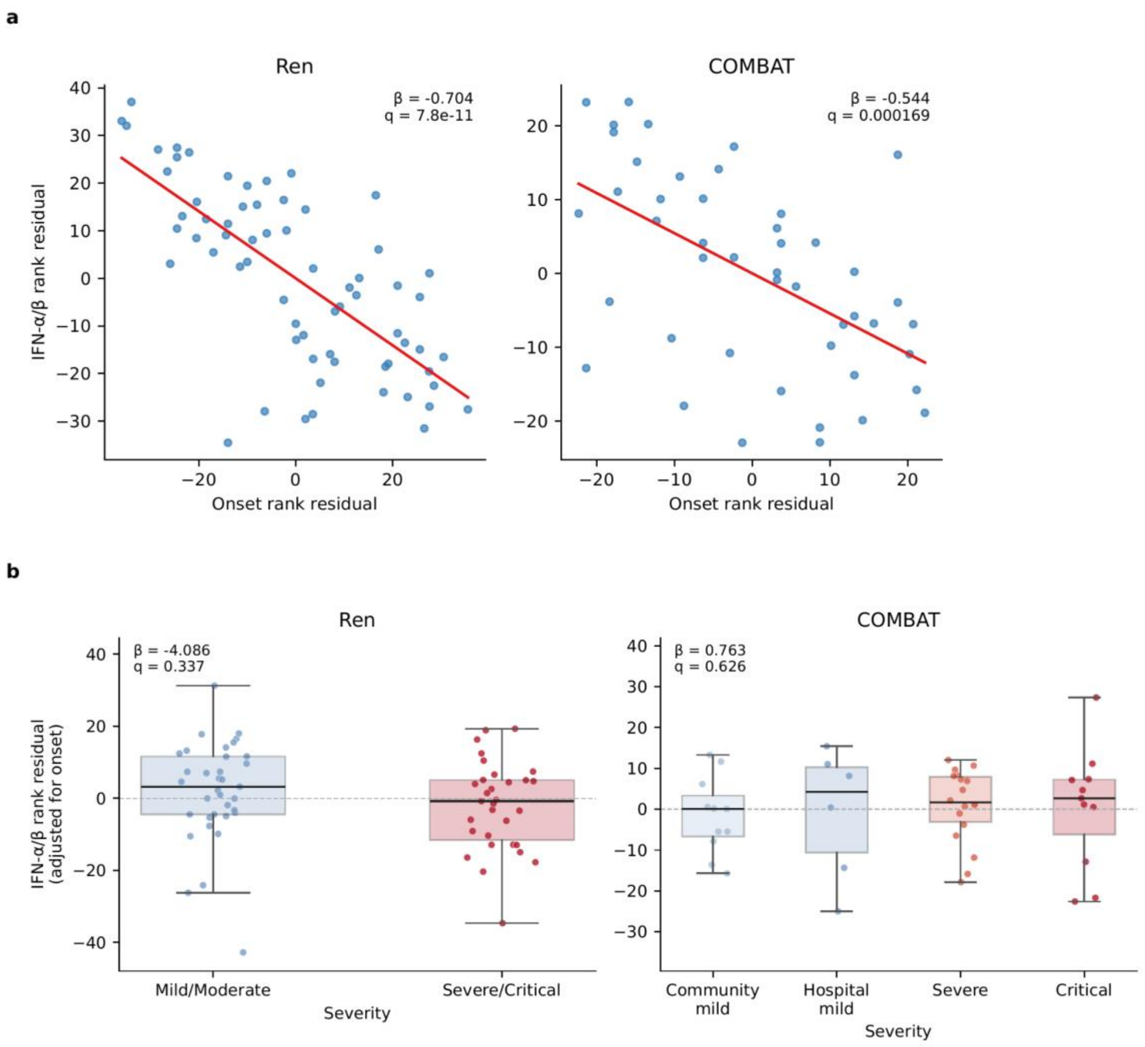


**Supplementary Fig. 7 | Interferon remodeling is associated with time since symptom onset but not disease severity in COVID-19.** (a) Association between the decoded between-patient IFN-α/β program and days since symptom onset after adjustment for disease severity, in Ren (n = 66) and COMBAT (n = 46). Axes show rank residuals from a rank-based linear model including both variables. Each point is a patient, and lines are least-squares fits. (b) Decoded between-patient IFN-α/β program by disease severity after adjustment for days since symptom onset. Severity is shown as two groups in Ren and as four ordered categories in COMBAT, following each cohort's clinical annotation. Each point is a patient, and boxes show medians and interquartile ranges. In both panels, the coefficient β from the rank-based model and the Benjamini-Hochberg-adjusted q are shown.

**Supplementary tables**

**Supplementary Table 1.** Dataset Characteristics, roles, and data sources of the single-cell and single-nucleus cohorts analyzed in this study.

| Dataset | Healthy donors (male/female) | Disease donors (male/female) | Cells | Age coverage | Accession |
|---|---|---|---|---|---|
| AIDA | 625 (275/350) | — | 1,265,624 | 19-77 | CELLxGENE ced320a1-29f3-47c1-a735-513c7084d508 |
| PsychAD (MSSM cohort) | 214 (132/82) | 212 (93/119) | 1,666,950 | 19-89 | CELLxGENE 37a17b78-4864-4a42-b67b-31c00962795a |
| Ren | 18 (16/2) | 92 (59/33) | 731,114 | 19-66 | GSE158055 |
| COMBAT | 10 (4/6) | 72 (32/40) | 378,408 | 18-79 | EGAS00001005493 |
| Stephenson | 22 (10/12) | 66 (38/28) | 369,970 | 20-69 | E-MTAB-10026 |

**Supplementary Table 2.** Ablation of endpoint-consistency and latent optimal-transport regularization in scCody-G and scCody-CT.

| Model | Validation Sinkhorn | Test Sinkhorn | Validation MAE | Test MAE | Validation Pearson r | Test Pearson r | Validation Source SD | Test Source SD |
|---|---|---|---|---|---|---|---|---|
| G: Full | **125.64** | **105.16** | 11.59 | 10.54 | 0.716 | 0.659 | 10.78 | 10.78 |
| G: VAE | 132.98 | 111.8 | 11.23 | 11.75 | 0.734 | 0.586 | 11.57 | 11.6 |

| | | | | | | | | |
|---|---|---|---|---|---|---|---|---|
| G:No endpoint | 126.4 | 105.36 | 13.19 | 10.59 | 0.591 | 0.688 | 11.23 | 11.47 |
| G: No latent OT | 125.9 | 105.2 | 12.01 | 10.45 | 0.744 | 0.672 | 10.84 | 10.92 |
| CT: Full | **128.38** | **105.48** | 11.09 | 10.89 | 0.682 | 0.63 | 11.4 | 11.6 |
| CT: VAE | 134.9 | 112.62 | 11.87 | 10.87 | 0.717 | 0.661 | 12.17 | 12.33 |
| CT: No endpoint | 128.97 | 106.47 | 11.46 | 10.94 | 0.599 | 0.659 | 11.12 | 10.55 |
| CT: No latent OT | 128.55 | 105.88 | 12.99 | 10.73 | 0.678 | 0.57 | 10.78 | 10.88 |

**Supplementary Table 3.** Internal age-holdout folds and selected checkpoints.

| fold | Internal holdout ages | External holdout ages | Source ages | Model | Selected epoch |
|---|---|---|---|---|---|
| 0 | 22,30,40,50,60,67 | 21,31,41,51,61,68 | 19,33,44,55,66 | scCody-G | 30 |
| | | | | scCody-CT | 29 |
| | | | | scNODE | 5 |
| | | | | scDiffEq | 500 |

| | | | | | |
|---|---|---|---|---|---|
| | | | | ARTEMIS | 19 |
| 1 | 23,32,42,52,62,66 | 21,31,41,51,61,68 | 19,33,44,55,67 | scCody-G | 24 |
| | | | | scCody-CT | 26 |
| | | | | scNODE | 6 |
| | | | | scDiffEq | 500 |
| | | | | ARTEMIS | 17 |
| 2 | 24,33,43,53,63,65 | 21,31,41,51,61,68 | 19,32,44,55,67 | scCody-G | 28 |
| | | | | scCody-CT | 30 |
| | | | | scNODE | 11 |
| | | | | scDiffEq | 475 |
| | | | | ARTEMIS | 16 |

**Supplementary Note 1 | Implementation of comparison generative dynamics models**

scNODE, scDiffEq, and ARTEMIS were trained on the same 2,000 genes, development donors, and three internal age-holdout fits as scCody (Methods). Hyperparameters for each method were tuned and checkpoints selected with the same internal age-holdout procedure as scCody. Search spaces and selected values are listed in Supplementary Table 3, and selected checkpoints in Supplementary Table 4. External held-out donors were used only for evaluation. Age localization, population generation, and differential-expression evaluation followed the procedures described for scCody. Each method computed Sinkhorn divergences in its own latent space, and cell-type labels for generated cells were assigned with a k-nearest-neighbour classifier trained on training cells. Age was rescaled to [0, 1] over 0–100 years.

The methods differed in how they simulate across age. scNODE integrates its ODE forward and backward in time, and ARTEMIS uses its learned forward and backward processes. scDiffEq learns only a forward-time stochastic process, so it was simulated forward from the youngest source population (19 years). All methods were evaluated over candidate ages 19–77.

*scNODE* was reimplemented following the original code, with the same model structure, ODE solver, and training losses: a variational autoencoder whose latent dynamics are an ODE with a ReLU drift network, integrated with the Euler method and trained with a Sinkhorn loss in expression space plus a latent-space term. To scale to the AIDA data, the autoencoder was pretrained on minibatches.

*scDiffEq* (v1.1.0) was used through its published interface, operating on 50 principal components with a potential drift and velocity-ratio regularization.

*ARTEMIS* was used through its published implementation, operating on a separately trained variational autoencoder, with age discretized into one-year steps.
